# Evolutionary diversification of the SymRK receptor family in land plants

**DOI:** 10.64898/2026.05.08.723708

**Authors:** Tora Fougner-Økland, Isaac Rodriguez-Arevalo, Athanasios Makris, Qichao Lian, Nadia Kamal, Korbinian Schneeberger, Martin Parniske, Martina K. Ried-Lasi, Katarzyna Parys

## Abstract

Plant receptor-like kinases (RLKs) are involved in diverse processes, ranging from growth and reproduction to interactions with microbes. Variation in the extracellular domains delineates several RLKs subfamilies, including the malectin-like domain leucine-rich repeat receptor-like kinases (MLD-LRR-RLKs). Symbiosis Receptor-like Kinase (SymRK) is the prototypical member of MLD-LRR-RLKs and is required for microbial accommodation in host roots during root endosymbiosis. Yet, comparative phylogenetic analysis of SymRK orthologs in the broader context of MLD-LRR-RLK subfamily evolution remains limited. In this study, we examined the inventory, phylogeny and clade-specific evolutionary and transcriptional characteristics of this receptor group. SymRK and its closest homologs are present in most land plant lineages and group into four major clades and six additional species-specific clades. These clades can be distinguished by their evolutionary characteristics as either conserved with reduced gene copy number changes (including SymRK) or expanded and diversified, as observed in clade IV. Clade IV dynamics are largely driven by tandem gene duplications, which often arise within gene clusters. We further analysed the evolutionary characteristics of MLD-LRR-RLKs at the population level in *Arabidopsis thaliana* accessions. We found that some genes are conserved across accessions and are therefore likely to be functionally important, whereas a subset of genes, often located within tandem clusters, are highly diverse and likely contribute to accession-specific adaptations. Finally, most *MLD-LRR-RLKs* in the *A. thaliana* Col-0 accession are expressed in roots and respond broadly to biotic stimuli at the transcriptional level. Notably, clustered genes frequently exhibited divergent expression profiles, suggesting transcriptional diversification. Together, we revealed two contrasting evolutionary characteristics among members of the MLD-LRR-RLK subfamily, potentially associated with their functions in plants.

## Background

Plants use receptor-like kinases (RLKs) to perceive diverse extracellular signals and translate them into appropriate cellular responses (Escocard de Azevedo Manhães et al., 2021; Shiu and Bleecker, 2001a). Numerous RLKs are encoded in plant genomes and orchestrate processes ranging from growth and development to the recognition of pathogenic and symbiotic microorganisms (Dievart et al., 2020).

RLKs share a conserved overall architecture comprising an extracellular domain (ECD), a single transmembrane helix and an intracellular kinase domain (Hohmann et al., 2017). Structural ECD diversity underlies the classification of RLKs into distinct sub-families (Shiu and Bleecker, 2003; Shiu and Bleecker, 2001b). Leucine-rich repeat (LRR)-RLKs represent the largest family in angiosperms (Shiu and Bleecker, 2001b). This receptor group has undergone extensive structural diversification, including variation in the number of repetitive LRR motifs as well as domain gains and losses, generating an expanded repertoire of structural variant types (Man et al., 2020). One subgroup is distinguished by the presence of an ancestral malectin-like domain (MLD), also found in *Catharanthus roseus* RLKs (*Cr*RLK1Ls) (Yang et al., 2021). Receptors containing an N-terminal MLD preceding a short LRR domain and an intracellular kinase domain are referred to as MLD-LRR-RLKs.

Symbiosis Receptor-like Kinase (SymRK) is the most extensively characterised member of the MLD-LRR-RLK subfamily. SymRK is indispensable for microbial uptake of *Glomeromycota* fungi in Arbuscular Mycorrhiza (AM) and nitrogen-fixing bacteria in Root Nodule Symbiosis (RNS) (Demchenko et al., 2004; Stracke et al., 2002). In addition to SymRK, the highly variable *Nodulation Specificity NS1* and *NS2* loci in *Medicago truncatula* encode MLD-LRR-RLKs that confer strain-specific compatibility in RNS (Liu et al., 2022; Yu et al., 2024). These examples support a role for MLD-LRR-RLKs in the regulation of symbiotic plant–microbe interactions.

Beyond roles in root endosymbiosis, functional characterisation of this receptor group has largely focused on *Arabidopsis thaliana*. *A. thaliana* has lost several important genes necessary for AM formation, including *SymRK* (Delaux et al., 2014). Instead, two close homologs, SymRK-Homologous Receptor Kinases 1 and 2 (SHRK1 and SHRK2), influence the accommodation of the obligate biotrophic oomycete and causal agent of downy mildew disease, *Hyaloperonospora arabidopsidis (Hpa)* (Ried et al., 2019). *A. thaliana* SHRK2, also known as Salt-Induced Malectin-like domain-containing Protein 1 (SIMP1), has also been linked to salt stress tolerance (He et al., 2021). Another well-characterised receptor, Impaired Oomycete Susceptibility 1 (IOS1), is encoded within a gene cluster of MLD-LRR-RLKs and contributes to susceptibility to *Hpa* and *Phytophthora parasitica* (Hok et al., 2014; Hok et al., 2011). IOS1 is also required for basal resistance against *Pseudomonas syringae,* indicating pathogen-dependent functions (Yeh et al., 2016). Moreover, *ios1* mutants exhibit enhanced sensitivity to abscisic acid (ABA) and salt stress (Cui et al., 2022; Hok et al., 2014), underscoring the complex functions of this receptor kinase. Allelic variants of Outgrowth-Associated protein Kinase (OAK), a member of a cluster of MLD-LRR-RLK in *A. thaliana*, are involved in accession-specific hybrid incompatibilities associated with aberrant phenotypes (Sageman-Furnas et al., 2022; Smith et al., 2011). Finally, the *Stress Induced Factor* (*SIF1-4*) genes are differentially regulated in response to both biotic and abiotic stresses (Yuan et al., 2018). For example, SIF2 promotes plant defence and flg22-induced stomatal closure (Chan et al., 2020), while SIF1 and SIF2 negatively regulate salt stress tolerance (Yuan et al., 2018). In addition to stress-related functions, MLD-LRR-RLKs such as Root Hair Specific (RHS) 6 and RHS16 are involved in root hair development (Won et al., 2009), and Canalization-related, Auxin-regulated Malectin-type RLK (CAMEL) is involved in auxin canalisation (Hajný et al., 2020). Collectively, these examples indicate that MLD-LRR-RLKs participate in diverse biological processes, including plant–microbe interactions, stress responses and development.

Phylogenetic inference of RLK-coding genes is often complicated by the extensive expansion and diversification of plant genomes, yet resolving the evolutionary history of a family of receptors can help to identify conserved receptor modules with core signalling function as well as rapidly evolving candidates that may contribute to species– or accession-specific adaptation to microbes (Fischer et al., 2016; Lehti-Shiu et al., 2009; Ngou et al., 2024). Among MLD-LRR-RLKs, SymRK has been extensively studied owing to its critical function in root endosymbiosis and its pronounced mutant phenotype, which facilitated its identification through forward genetic screens in legumes (Endre et al., 2002; Stracke et al., 2002). However, the evolutionary characteristics of SymRK orthologs in relation to the broader MLD-LRR-RLK subfamily remain incompletely resolved. Previous studies either focused on SymRK in the context of common symbiosis genes (Radhakrishnan et al., 2020) or grouped MLD-LRR-RLK members into the LRR-I subgroup and examined them within the broader context of the LRR-RLK superfamily (Fischer et al., 2016; Kileeg et al., 2023; Man et al., 2020). In this work we employed a refined strategy and investigated the phylogenetic trajectories of MLD-LRR-RLKs in land plants, using SymRK as the founding member. We inferred clade-specific characteristics of this receptor subfamily, compiled a receptor inventory and used publicly available transcriptomic resources to assess expression patterns in *A. thaliana*.

## Results

### Members of the SymRK receptor family cluster into four major clades in land plants

To investigate the evolutionary trajectories of the *SymRK* receptor family and to resolve orthology-paralogy relationships, we performed a phylogenetic analysis of 46 published plant genomes (**Fig. S1, Table S1**). Where possible, we restricted our dataset to species representing major land plant lineages with high-quality diploid genome assemblies and minimal recent whole genome duplications (**Table S1**). Orthogroups were inferred across these genomes using OrthoFinder (Emms and Kelly, 2019). This returned a total of 784 protein-coding genes annotated as homologs of the SymRK receptor family. In addition, we performed an unbiased domain-based search for MLD-LRR-RLKs and identified 100 additional proteins not captured by the OrthoFinder approach. Among these candidates, 73 sequences belonged to a clade with higher similarity to the *Cr*RLK1L family and the detected LRRs overlapped with other well-annotated domains. Because these sequences lacked clear phylogenetic or domain-architecture homology to MLD-LRR-RLKs, we excluded this clade from our analysis (**Fig S2b**). The unsupervised phylogenetic analysis of all identified proteins revealed spurious long branches, which were subsequently closely examined (**Fig. S2b**). Some of the identified gene models consisted of short gene fragments that did not encode full-length domains (<250 residues), whereas others encoded multiple MLDs or kinase domains in tandem (>1200 residues). Based on these observations, we filtered proteins by length and removed those that did not start with an initiating methionine. Finally, our dataset comprised 694 high-confidence protein-coding gene models of putative SymRK-related MLD-LRR-RLKs (**Fig. 1** and **Fig. S2**).

**Figure 1:**
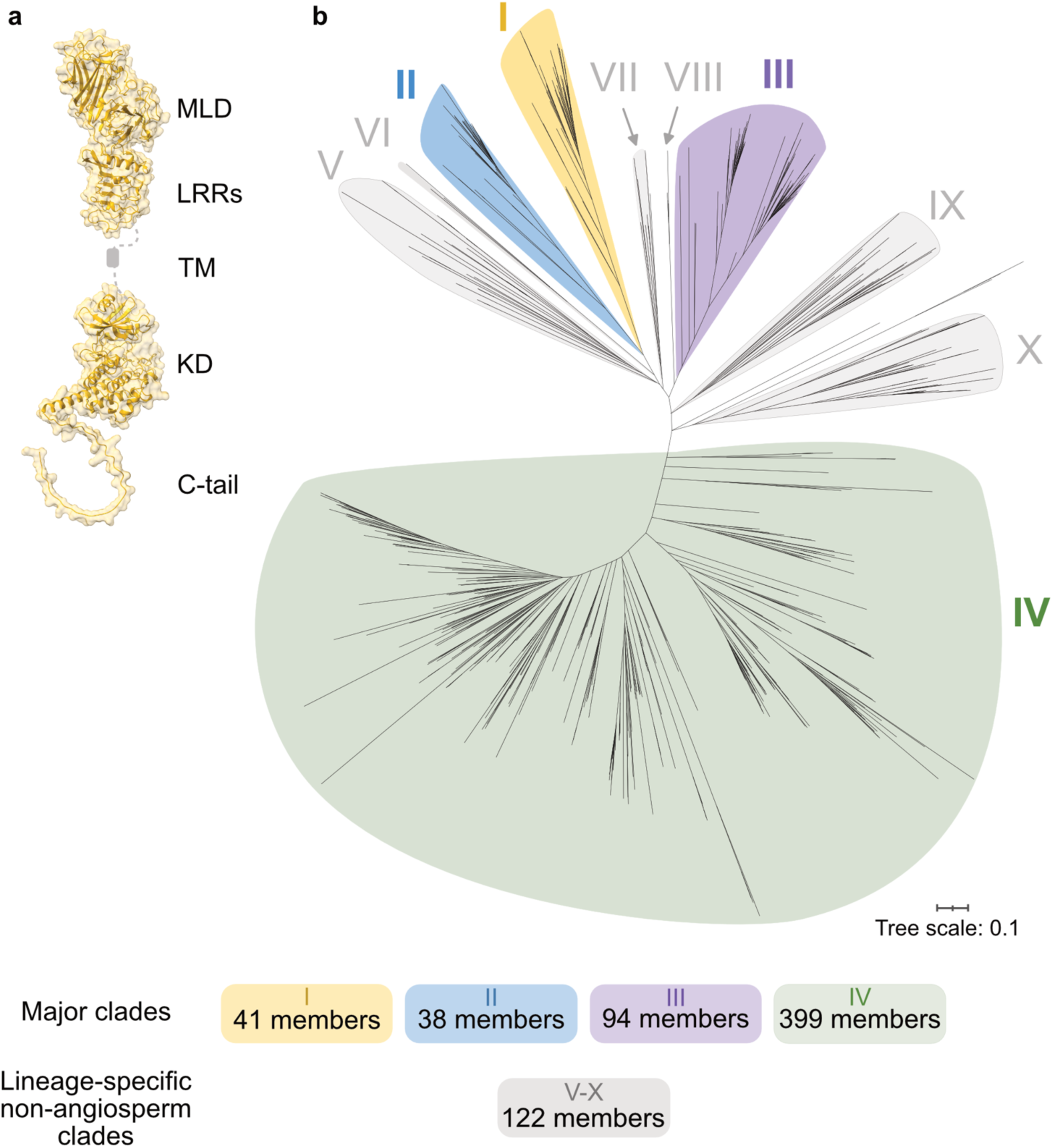
SymRK receptor family members form four major clades in land plants. **a** – AlphaFold 3 structural prediction of *Lotus japonicus* SymRK, highlighting the multi-domain nature of this receptor family. The transmembrane domain and the adjacent unstructured juxtamembrane regions are depicted schematically. MLD – Malectin-like Domain, LRR – Leucine-rich Repeats, TM, transmembrane domain, KD, kinase domain. **b**– Unrooted maximum-likelihood phylogenetic tree of proteins homologous to SymRK. Four major clades are resolved: clade I (yellow), clade II (blue), clade III (purple) and clade IV (green). Clades V – X, which represent lineage-specific expansions in non-angiosperm species are highlighted in grey. Clade V – *Marchantia,* Clade VI – *Selaginella*, Clade VII – *Physcomitrella*, Clade VIII – *Ginkgo*, Clade IX – *Anthoceros*, Clade X – *Azolla, Ceratopteris*. Bootstrap and orthogroup support for clade definitions are shown in Fig. S3. Below the tree, the clade-wise phylogenetic placement of SymRK receptor homologs is shown. Note that the number of protein-coding genes assigned to each clade is unequal. The tree was inferred from high-confidence protein sequences identified across 46 plant genomes. Sequences were aligned with MAFFT, trimmed using BMGE and tree built using RAXML-NG.

A maximum likelihood tree of the SymRK-homologous protein sequences resolved ten clades of unequal size in land plants (**Fig. 1**). We overlaid taxonomic information onto the phylogenetic tree and found that angiosperm homologs of SymRK clustered consistently in four clades, which aligned with the orthogroup assignments (**Fig. S3**). Based on this pattern, we designated four major clades (I-IV) and additional lineage-specific clades (V-X) in non-angiosperms (**Fig. 1**, **Fig. S3**). We analysed the placement of eight *L. japonicus* genes encoding MLD-LRR-RLKs, referred to as SymRK and SHRKs, across the established clades and found them distributed among the four major clades I-IV (**Fig. S4**). Notably, SymRK was confined to clade I, which contained only SymRK orthologs (**Fig. S5a**). Clades I and II each formed well-supported groups with bootstrap values >97 (**Fig. S3**). These two clades shared a distinct branch that separates them from all other MLD-LRR-RLKs (**Fig. 1**), consistent with an early duplication during land plant evolution. Support for clades III and IV was weaker, and the clades were delineated based on strong bootstrap support within angiosperm homologs as well as monophyletic clustering of orthogroup members. Based on this delineation, members of clades I-III were identified in bryophytes (*Physcomitrella patens*, *Marchantia paleacea*, *Marchantia polymorpha*) and lycophytes (*Selaginella moellendorffii*), suggesting that the diversification within the SymRK receptor family occurred early in land plant evolution, before the divergence of bryophyte and vascular plant lineages.

In addition to the major clades, several species harboured MLD-LRR-RLKs that grouped within clades V-X (**Fig. 1**). These lineage-specific clades were dominated by specific species, including *Ceratopteris richardii*, *M. polymorpha*, *M. paleacea* and *P. patens* and may represent independent lineage-specific expansion events. Taken together, our analyses identified four major clades within the SymRK receptor family in land plants, together with six lineage-specific non-angiosperm clades, each containing different numbers of members.

### Contrasting patterns of expansion and retention among major clades of the SymRK receptor family

While clade assignment and orthogroup inference confirmed that SymRK receptor family members are broadly present across land plant lineages, notable inter-clade differences were observed in gene copy numbers and receptor structural variants (**Fig. 1**). To address this, we analysed presence-absence patterns and gene copy number variation across each clade (**Fig. 2**). Clade I members, or SymRK orthologs, were consistently detected in species that engage in AM (e.g. *M. truncatula*, *M. paleacea* and *Oryza sativa*) or orchid mycorrhiza (*Phalaenopsis equestris, Vanilla planifolia*), but were absent from species that have lost mycorrhizal symbiosis, such as *A. thaliana* and *Beta vulgaris* (**Fig. 2**, **Fig. S5a**). This distribution corroborates previous findings (Delaux et al., 2014). An apparent exception was *Acorus gramineus*, where no *bona fide* SymRK ortholog was identified. Instead, BLAST searches revealed an LRR-RLK lacking both the MLD and a core part of the kinase, including the activation loop, which is predicted to impair kinase function (**Fig. S5b**). In most genomes, a single *SymRK* gene was maintained. Exceptions included *Helianthus annuus, P. equestris* and *V. planifolia*, possibly due to recent whole genome duplication or partial reduplication (**Fig. 1** and **Fig. S5**) (Badouin et al., 2017; Piet et al., 2022). More than one gene was also identified in *Daucus carota*, which harboured one full-length copy and a second truncated variant that carried a deletion in the C-terminal kinase lobe, resulting in a protein lacking critical functional domains (**Fig. S5b**). These observations highlight a tendency toward maintaining a single seemingly functional *SymRK* copy per genome.

**Figure 2:**
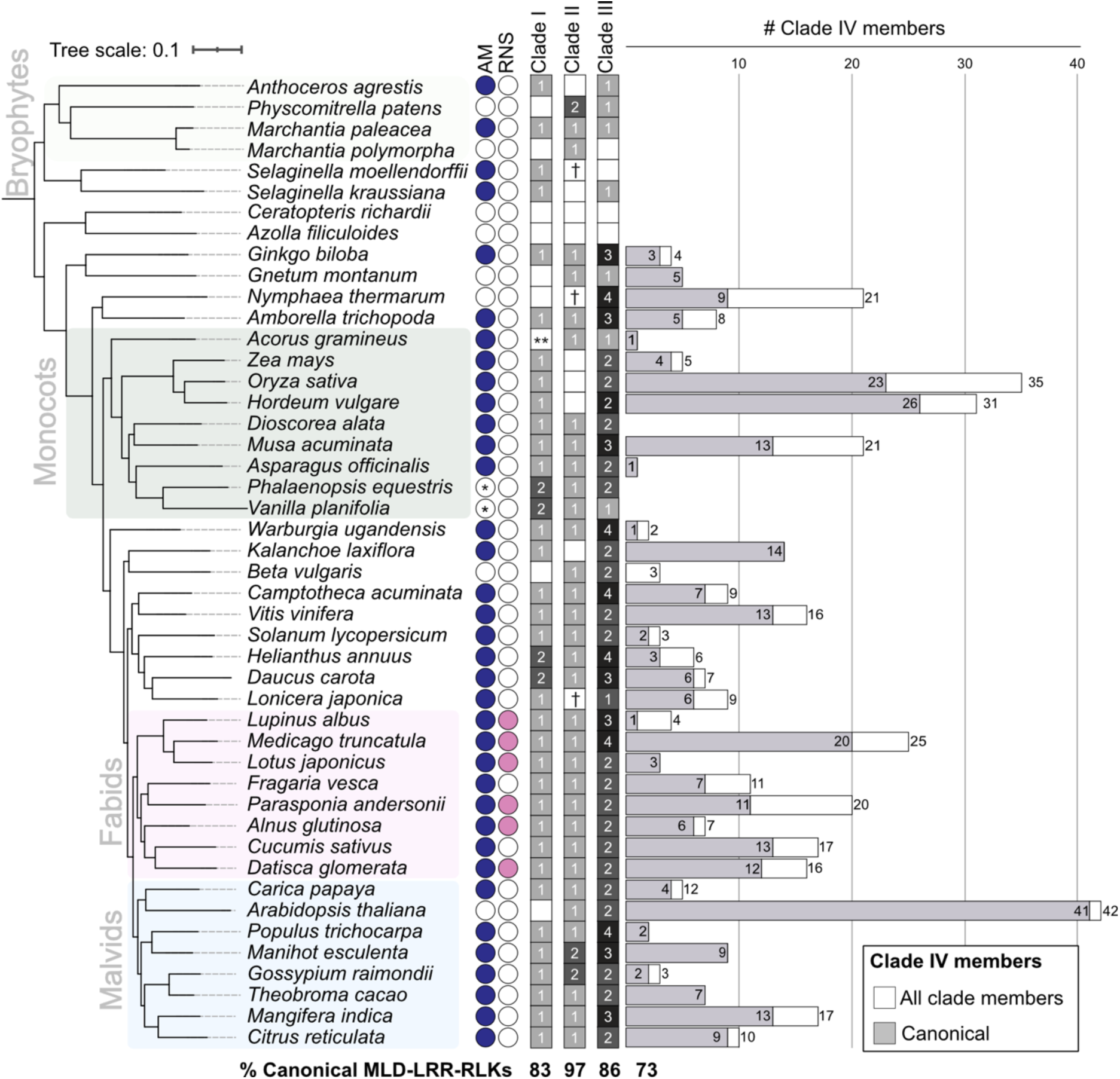
SymRK receptor family members are prevalent in land plants and display clade-specific phylogenetic characteristics. For each major clade (I-IV), the grid shows the presence, absence and number of identified receptor family members across the selected plant species. In the largest clade (clade IV), stacked white bars indicate the total number of clade members per species, while grey bars represent the subset of members classified as canonical MLD-LRR-RLKs (containing MLD, LRR, TM, and kinase domains). AM – Arbuscular Mycorrhiza; RNS – Root Nodule Symbiosis. * – *Phalaenopsis equestris* and *Vanilla planifolia* form orchid mycorrhiza but not AM. ** – No ortholog was identified in *Acorus gramineus*, but a potential ortholog with functionally critical deletions was identified using BLAST (see also Supplementary Figure 5). † – A single homolog was identified, but removed during filtering by size constraints (250-1200 aa).

Clade II displayed a conservation pattern similar to that of clade I, exemplified by predominantly a single gene copy per genome, except in species with recent genome duplications, such as *H. annuus* and *P. equestris* (**Fig. 2**). For simplicity, we refer to members of this clade as SHRK3 orthologs (**Fig. S6**). We observed that *SHRK3* orthologs are absent from aquatic ferns (*Azolla filiculoides* and *C. richardii*) and from Poaceae (*Zea mays, O. sativa* and *Hordeum vulgare*) (**Fig. 2**, **Fig. S6**). We interpret this pattern as consistent with lineage-specific losses that are not obviously associated with root endosymbiotic status, and therefore distinct from what we observed for *SymRK*. Interestingly, within our dataset, no species was found to simultaneously carry multiple copies of both *SymRK* and *SHRK3* (**Fig. 2**), suggesting that concurrent expansion of these two clades is uncommon among the sampled species, although additional sampling may reveal exceptions.

No bryophyte or lycophyte species carried more than one gene within Clade III (**Fig. 2** and **Fig. S7**). In contrast, in seed plants, Clade III split into two subclades, III-A and III-B, likely resulting from a duplication event predating the divergence of gymnosperms and angiosperms (**Fig. S7**). These correspond to *SHRK1* and *SHRK2*/*BSR050/SIMP2,* previously described in *A. thaliana* (He et al., 2021; Ried et al., 2019; Xu et al., 2014). Angiosperms generally carried more copies of clade III genes than clade I or II, although copy numbers did not exceed four per genome.

Gene trees for clades I-III were largely congruent with species trees of angiosperms (**Fig. S5-S7**), suggesting predominantly speciation-driven divergence. By contrast, clade IV showed striking fluctuations in copy number across angiosperm families (**Fig. 2**). In particular, large expansions were observed in monocots (*Musa acuminata, O. sativa, Hordeum vulgare)*, in *Nymphaea thermarum* and in several rosids (*M. truncatula, A. thaliana)* (**Fig. S8**). These expansions were not obviously attributable to broad taxonomic relationships or a clear ecological pattern, suggesting that they may have occurred independently in different lineages. Notably, clade IV contains many MLD-LRR-RLKs for which corresponding mutants have been characterised in *A. thaliana* and in *M. truncatula* (**Fig. S8**). These taxonomically inconsistent copy number changes indicate that clade IV has undergone a complex evolutionary trajectory, shaped by lineage-specific expansions and possibly also by diversification at the population level.

### Comparative protein sequence analysis reveals distinct characteristics of the major clades of the SymRK receptor family

The contrasting patterns of gene presence/absence and copy number variation across clades I-IV suggested that these clades may have evolved under different selective constraints. To address this, we compared diversity within clades at the domain and residue level (**Fig. S9** and **S10**).

We assessed the overall degree of structural variability among members of each clade using Pfam domain searches and PhytoLRR to profile variation in LRR array length (**Fig. S9**). Based on these analyses, we defined two categories of protein variants: “canonical”, which contain the MLD, LRR, TM and kinase domain, and “non-canonical”, which lack at least one of these domains. Because domain prediction approaches fail to capture variability in the unstructured juxtamembrane and C-terminal regions, which lack conserved motifs and are prone to length variation, we also examined intra– and extracellular domain lengths (**Fig. S9a**). Clades I-III retained a canonical MLD-LRR-RLK domain architecture in more than 83% of sequences (**Fig. 2**, **Fig. S5a, Fig. S6-S8**). A small cluster of sequences with shorter ectodomains was observed in clade I, which primarily accounts for the non-canonical variants in this clade (**Fig. S9a**). This pattern can be attributed to lineage-specific MLD losses in the analysed species of commelinids (*M. acuminata, H. vulgare, O. sativa, Z. mays)* (**Fig. S5a**), consistent with previous reports of domain losses in monocots (Miyata et al., 2023). In addition, clade I members had the fewest LRRs, in some cases only two, whereas more than 70% of clade II and III members had four or more LRRs (**Fig. S9b**). Clade II members showed particularly low structural variation, characterised by a predominant canonical architecture and the highest number and frequency of LRRs among the analysed clades (**Fig. S9a, b**). These ectodomain variations may reflect clade-specific constraints on ectodomain architecture.

To determine whether the observed clade-specific structural ectodomain variability was also reflected at the sequence level, we analysed a conserved signature motif of the ectodomain. The GDPC (glycine-aspartic acid-proline-cysteine) amino acid sequence, with flanking residues, first described in *L. japonicus* SymRK, is situated between the MLD and LRRs in the ectodomain of MLD-LRR-RLKs and has been linked to MLD release from the receptor (Antolín-Llovera et al., 2014; Chakrabarti et al., 2026; Kileeg et al., 2023). We asked if the motif and its surrounding region displayed clade-specific signatures. Alignment of this region across clade members revealed a shared consensus sequence: WxxDPCxPxxWxxxxC, where W represents tryptophan and x any amino acid (**Fig. S9c**). Despite this overall conservation, clade-specific signatures emerged. In particular, the GDPC motif was altered in clade II to DDPC. Clade III featured a second W at position 17 in the sequence logo, which was absent from the other clades. Furthermore, while a conserved P at position +4 relative to xDPC was present in clades I-III, it was missing in clade IV, which instead displayed an aromatic residue, tyrosine (Y) or phenylalanine (F), at position 12 in the logo. In summary, the GDPC motif and its flanking region serve as a clade-specific fingerprint within this receptor family.

While clades I-III exhibited comparatively limited structural variability, mainly affecting ectodomain architecture, clade IV displayed the highest degree of overall structural variation (**Fig. S9a**). This clade contained a higher proportion of non-canonical structural variants, representing 27% of all members (**Fig. 2**). These variants were considered non-canonical either because they lacked one or more domains or domain features (the MLD, individual LRR motifs, the entire LRR domain, or the kinase domain), or because they carried additional domains with no homology to MLD, LRRs or kinase domain (**Fig. S8, S9a**). The frequency of such events was not obviously correlated with broad taxonomic position, suggesting that structural diversification in clade IV occurred repeatedly in different lineages rather than reflecting a single shared ancestral event. Variation in domain length also arose from expansion and contraction in unstructured regions, particularly the juxtamembrane segment or C-terminal tail. Overall, these observations indicate higher structural plasticity in clade IV than in the more constrained architectures of clades I-III.

To further evaluate polymorphism levels among members of the major clades, we conducted amino acid sequence identity analyses. Clades I-III exhibited higher overall amino acid conservation than clade IV, and no significant differences were observed among clades I-III (**Fig. S10a**). To assess whether extracellular and intracellular domains evolved under different constraints, we compared pairwise sequence identity of MLDs and kinase domains across the major clades (**Fig. S10b**). We observed higher amino acid identity for the kinase domains compared to the MLDs across all clades. This asymmetric pattern is in line with previous analyses of positive and negative selection in LRR-RLKs (Man et al., 2023). Notably, MLDs of clade II and III members displayed higher sequence identity than those of clade I and IV members. This tendency did not apply to kinase domains, where all clades, with the exception of clade IV, displayed similarly high levels of conservation (**Fig. S10**).

In summary, our analyses revealed two distinct evolutionary dynamics within MLD-LRR-RLKs: (i) a conserved trajectory for clade I-III, characterised by sequence conservation and predominantly speciation-associated divergence, and (ii) a more dynamic trajectory for clade IV, reflected at both the sequence and domain-architecture levels.

### MLD-LRR-RLKs expansions in clade IV are associated with tandem gene duplication events

The observed variation in the number of paralogs within clades IV-X prompted us to investigate the genomic arrangements associated with expansion and diversification of these members. Tandem gene duplications have been reported to contribute to the expansion of receptor kinase families, including LRR subfamily I in *A. thaliana*, which includes MLD-LRR-RLKs (Kileeg et al., 2023). To determine whether expanded clades IV-X were associated with events of tandem duplications, we analysed the genomic position of each gene and its gene neighbourhood within each species. Genes were classified according to their genomic arrangement as: i) single dispersed paralogs, if no other MLD-LRR-RLK-encoding genes were located in their proximity, ii) proximal paralogs, separated by no more than two intervening genes, iii) tandem paralogs, located immediately adjacent to another MLD-LRR-RLK, or iv) tandem clusters, defined as arrays of more than two paralogs within a single genomic block (**Fig. 11a**).

We found that clades V-VIII were mostly composed of paralogs present as single dispersed copies. In clades IX and X, 50% of the paralogs occurred either as proximal or tandem paralogs, or as part of tandem arrays (**Fig. S11b**). By contrast, a large proportion of clade IV MLD-LRR-RLKs were encoded within tandem gene arrays (**Fig. 3a**). The frequency of tandem arrays varied among the species analysed and did not simply follow species’ phylogenetic relationships (**Fig. 2**, **Fig. 3a**). However, we observed a positive correlation between the total number of clade IV MLD-LRR-RLKs in a given species and the number of these genes located in tandem arrays (**Fig. S11c**). No duplication events of MLD-LRR-RKs were detected in *Gnetum montanum*, *Acorus gramineus*, *Asparagus officinalis*, *Solanum lycopersicum* and *Gossypium raimondii*; notably, each of these species encoded fewer than five clade IV MLD-LRR-RLKs. In summary, our findings suggest that the expansion of clade IV is primarily due to tandem gene duplications.

**Figure 3:**
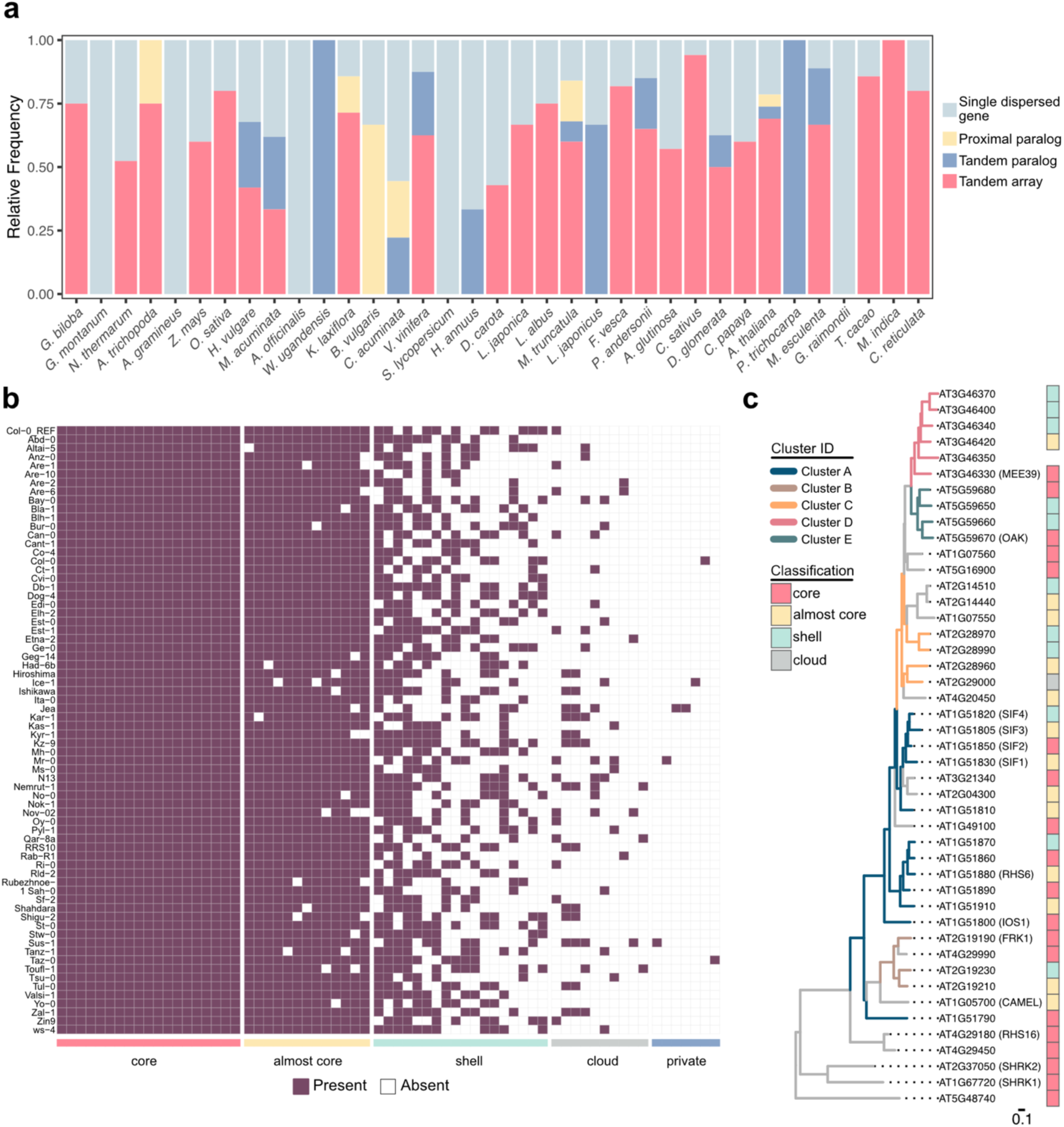
Clustered genes drive the uneven expansion of clade IV members of the SymRK receptor family. **a**– Chromosomal arrangement of clade IV genes in each species organised by species tree as in Fig. 2, represented as relative frequencies of genes occurring either as singletons or with additional nearby copies. Genes with nearby copies are classified as proximal duplicates, tandem duplicates, or gene clusters within genomic regions analysed for each species. **b –** Copy number fluctuations in Clade IV are observed at the population level. The heat map shows the presence–absence of clade IV orthologous genes across 69 *A. thaliana* accessions. Based on frequency of occurrence across the accessions, the genes are classified as core (present in all accessions), almost core (present in at least 62 accessions), shell (found in at least 13 accessions), cloud (occurring in at least 2 accessions), and private (unique to one accession). **c –** Phylogenetic tree for all *A. thaliana* MLD-LRR-RLKs, including clade I (AT5G48740) and clade II (SHRK1 and SHRK2) members. Cluster annotation is displayed with a different colour of the branches, while presence-absence polymorphism across *A. thaliana* accessions is highlighted with coloured squares. Common names are indicated.

We asked whether the dynamic gene evolution detected in clade IV could also be observed at the population level. *A. thaliana* encodes the highest number of MLD-LRR-RLKs in our dataset, with 42 clade IV members, of which 66% form tandem arrays (**Fig. 2**, **Fig. 3a**). To address this, we identified all MLD-LRR-RLK-encoding genes in 69 *A. thaliana* accessions with chromosome-level assembly (Lian et al., 2024). *A. thaliana* accessions encode variable numbers of MLD-LRR-RLKs, ranging from 36 in Rubezhnoe to 47 in Kz-9 (**Fig. 3b**, **Fig. S11d**). The population-level gene copy variation pointed to recurrent gene gains and losses within tandemly arrayed regions (**Fig. 3c**, **Fig. S12-S14**).

To understand the variation in MLD-LRR-RLKs across ecotypes, we clustered them into orthogroups (OG) based on sequence similarity. The OGs were classified into five pangenome categories according to how often each OG occurs across the accessions: core (present in all accessions), almost core (present in at least 62 accessions), shell (found in at least 13 accessions), cloud (occurring in at least 2-12 accessions), and private (unique to one accession) (**Fig. 3b**). Among the identified MLD-LRR-RLKs, 19 (28%) constitute the core set, while 13 (19%), 18 (26%), 10 (15%), and 7 (10%) represented the almost core, shell, cloud, and private groups, respectively (**Fig. 3c**). Approximately half of the MLD-LRR-RLKs are maintained within the core and almost-core categories, consistent with a stronger evolutionary constraint and perhaps functional importance, whereas shell, cloud and private groups captured variable components of the MLD-LRR-RLK repertoire.

By integrating pangenome classification with phylogenetic analysis of MLD-LRR-RLKs in *A. thaliana* Col-0, we observed that almost all cloud and shell genes were part of tandem gene arrays (**Fig. 3c**). Thus, tandem arrays represent major contributors to *MLD-LRR-RLK* copy-number variation across *A. thaliana* accessions. In the opposite fashion, core and almost core genes tended to occur as single dispersed genes and were less frequently associated with gene duplications. This split reinforces the idea that much of the variation in gene family size among MLD-LRR-RLKs arises from tandem arrays, suggesting that these genomic regions could serve as sources of genetic variation.

### Transcriptional profiling of clustered genes in *Arabidopsis thaliana* revealed heterogeneous expression patterns

To assess the overall expression profile of *A. thaliana MLD-LRR-RLKs* and to evaluate whether clustered genes show diverging expression patterns, we performed an exploratory meta-transcriptomic analysis using a pre-compiled RNA-sequencing database (Yu et al., 2022). The database compiles published *A. thaliana* transcriptomes, including tissue-specific expression and transcriptional responses to biotic stimuli. Only libraries with comprehensive experimental descriptions in the Col-0 background were analysed. After preliminary quality assessment, we obtained a final set of 878 libraries for tissue expression and 372 for biotic stress responses.

Analysis of tissue-specific expression revealed distinct patterns among the clades. The conserved clade II gene (*AT5G48740*) and clade III genes (*SHRK1* and *SHRK2*) were expressed across multiple plant tissues (**Fig. 4**). A similar pattern was observed for *IOS1*, while a small subset of genes showed preferential expression in aerial tissues, including *SIF3*, *OAK*, *SIF4*, *AT3G46370* and *Flg22-induced Receptor-like Kinase 1 (FRK1)*. By contrast, many clade IV genes were preferentially expressed in roots and seedlings (**Fig. 4**). Nine genes had very low or undetectable transcript abundance in the analysed datasets (**Fig. S16-17**), which may indicate pseudogenisation, low basal expression or highly restricted expression in specialised cell types or conditions not represented in the available datasets. This analysis, based on multiple independent libraries, was broadly consistent with the tissue expression data from (Klepikova et al., 2016) (**Fig. S15** and **Fig. S16**). We found no clear associations between tissue expression patterns and physical gene cluster identity (**Fig. 4**). While some paralog clusters, in particular cluster C and E, which includes *OAK,* displayed similar expression patterns across tissues, most clusters contained genes with distinct expression profiles (**Fig. 4**).

**Figure 4.**
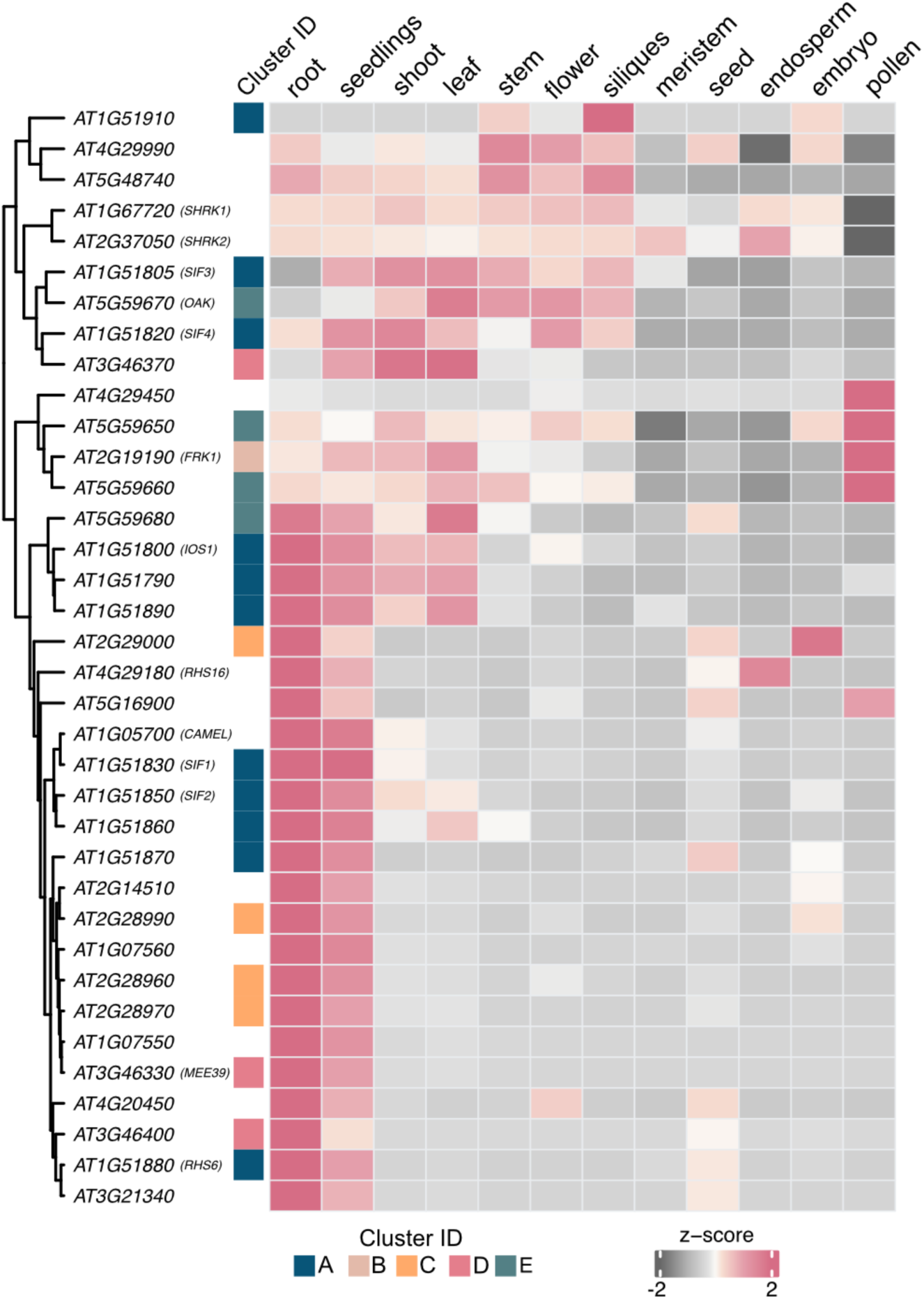
Most *MLD-LRR-RLKs* in *Arabidopsis thaliana* are expressed predominantly in the root. The heat map displays normalised gene counts for *A. thaliana* MLD-LRR-RLKs across various tissues, which are listed column-wise at the bottom of the heat map. Transcript abundance is represented by shades of pink and grey. Genes were hierarchically clustered based on similarities in their co-expression profiles. Gene IDs are shown on the left of the heat map, together with common names where available. Cluster assignment is indicated by coloured squares adjacent to the gene IDs, with each colour denoting a distinct cluster.

Several *MLD-LRR-RLKs*, such as *FRK1*, *IOS1*, *SHRK1* and *SHRK2,* have previously been associated with a role in biotic interactions (Asai et al., 2002; Hok et al., 2014; Hok et al., 2011; Ried et al., 2019). We therefore asked whether transcriptional responsiveness to biotic stimuli might represent a shared feature of this family of receptors and if the dynamic evolution of tandem arrays could be linked to plant-biotic interactions. To investigate this, we analysed expression profiles of all *MLD-LRR-RLK* genes in *A. thaliana* in response to various biotic agents (**Fig. 5**). Transcriptional changes were examined across three categories: shoots, roots, and seedlings. This approach allows accounting for tissue-specific expression of *MLD-LRR-RLK* genes and the primary sites of infection or interaction. Many *MLD-LRR-RLKs* displayed transcriptional changes in response to at least one biotic condition (**Fig. 5**). However, at this level, we did not observe clear organism-specific response patterns across the receptor family (**Fig. 5**), and several *MLD-LRR-RLKs* showed weak or no detectable transcriptional responses under the analysed conditions. (**Fig. S17**).

**Figure 5.**
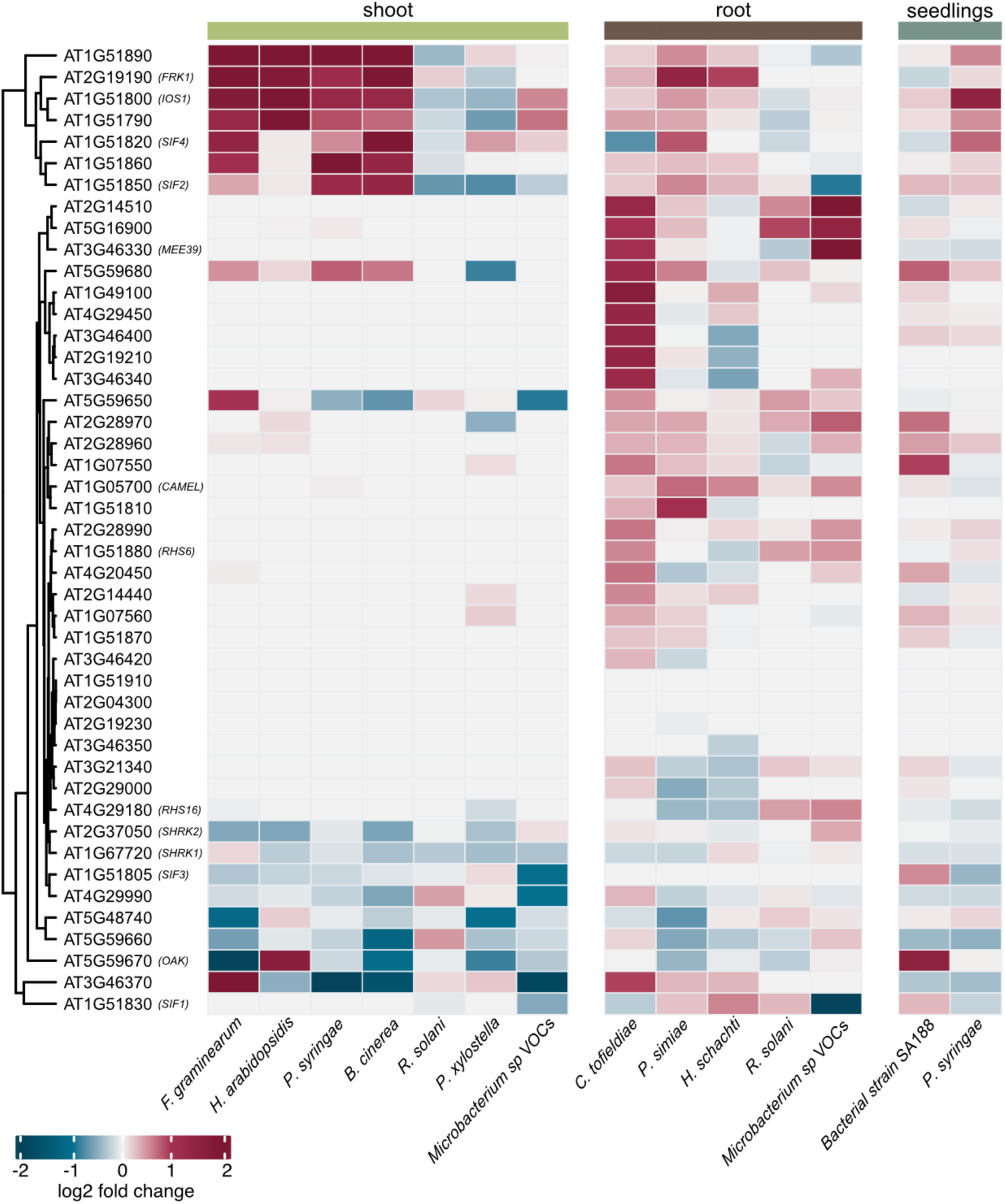
MLD-LRR-RLKs in *Arabidopsis thaliana* display transcriptional diversification in response to biotic stresses. The heat map depicts the weighted by precision log2 fold changes of *A. thaliana MLD-LRR-RLKs* in response to different biotic agents, which are indicated column-wise at the bottom of the heat map (*Fusarium graminearum, Hyaloperonospora arabidopsidis, Pseudomonas syringae, Botrytis cinerea, Rhizoctonia solani*, *Plutella xylostella, Colletotrichum tofieldiae, Pseudomonas simiae, Heterodera schachtii*). Normalised gene counts across 372 libraries (see Methods) are presented, with transcript abundance represented by shades of blue and burgundy. Genes were hierarchically clustered based on similarities in their expression profiles. Gene IDs are shown on the left of the heat map, accompanied by common names when available. VOCs: Volatile organic compounds.

Hierarchical clustering of co-expression patterns in shoots revealed two gene subgroups that exhibit strong induction or repression in response to several microbes, including the necrotrophic fungi *Fusarium graminearum* and *Botrytis cinerea*, the biotrophic oomycete *Hpa* and the hemibiotrophic bacterium *Pseudomonas syringae* (**Fig. 5**). The subgroup of strongly induced genes included previously described receptors involved in response to microbes, including the immune response marker gene *FRK1* (Asai et al., 2002). These genes also exhibited strong responses to microbial molecular patterns (**Fig. S18**) in a separate elicitor dataset (Bjornson et al., 2021). This suggests that they generally safeguard plant basal defence responses in the shoot while also exhibiting low-level induction by root-associated microbes, such as the biotrophic fungi *Colletotrichum tofieldiae* and *Pseudomonas simiae*. Notably, except for *FRK1*, many of the *MLD-LRR-RLKs* that respond to microbes were part of the tandem array cluster A on chromosome 1, which is conserved across *A. thaliana* accessions (**Fig. 3c**). This pattern suggests a co-regulation of cluster A genes, possibly due to their genomic proximity. A second subgroup, including *SHRK1, SHRK2, SIF3* and *OAK*, showed predominantly reduced transcript abundance under several biotic conditions, reflecting diversified transcriptional responses among *MLD-LRR-RLKs*. In roots, many *MLD-LRR-RLKs* were induced, a trend that mirrors the one present in seedlings (**Fig. 5**). This broader transcriptional induction in roots may be associated with the specific expression of many *MLD-LRR-RLKs* in root tissues (**Fig. 4**).

In summary, transcriptional profiling of *MLD-LRR-RLKs* under diverse biotic stresses suggests that some members of this gene family are transcriptionally integrated into biotic-stimulus response programmes. However, no clear associations could be established between microbial species, lifestyles or individual biotic agents and the corresponding gene response patterns. Notably, the tendencies shown in **Fig. 5** should be interpreted with caution, as this meta-analysis captures broad trends that may vary across time points, developmental stages or experimental conditions.

## Discussion

### Opposite evolutionary traits detected in the SymRK receptor family

Our phylogenetic framework classified the SymRK receptor family into four major clades that capture the principal diversity of MLD-LRR-RLKs in angiosperms, and six additional lineage-specific clades in non-flowering plants (**Fig. 1**). Clades I-III are characterised by low copy numbers, in contrast to the expanded clade IV. Similar clade-specific characteristics have been described for other RLKs, suggesting that asymmetric selective pressures can shape the evolutionary dynamics of receptor families (Hosseini et al., 2020; Kileeg et al., 2023; Shiu and Bleecker, 2003; Vaattovaara et al., 2019). As a result, phylogenetic patterns are frequently used to infer putative functional predispositions (Man et al., 2023). Consistent with this framework, several features observed for clade I-III members are compatible with conserved signalling functions. By contrast, the expansion, structural diversity and transcriptional responsiveness of clade IV suggest that members of this clade are more likely to be involved in plant-microbe interactions, equipping plants with genetic tools that could contribute to adaptive plasticity in response to biotic interactions.

### Members of clades I and II exhibit patterns consistent with dosage sensitivity

Clade I (SymRK) and clade II (SHRK3) members are predominantly retained as single-copy genes across the analysed plant genomes, suggesting that their functions may be dosage-sensitive. Such retention patterns are commonly observed for tightly regulated genes involved in cellular homeostasis and organismal fitness (De Smet et al., 2013). The single-copy status of *SymRK* may result from its proposed role in dampening flg22-triggered immune responses during rhizobial accommodation (Feng et al., 2021). The latter occurs through inhibition of the kinase activity of Brassinosteroid Insensitive 1 Associated Kinase 1 (BAK1), a positive regulator of defence signalling (DeFalco and Zipfel, 2021). Accordingly, ectopic expression of SymRK reduces flg22 sensitivity, suggesting that elevated SymRK levels may compromise immune responsiveness under certain conditions (Feng et al., 2021). This may underlie the absence of *SymRK* in species that have lost AM (Delaux et al., 2014; Radhakrishnan et al., 2020) and the potential intermediate case of gene erosion, as observed in *Acorus*, where *SymRK* key functional regions appear to be lost (**Fig. S5b**).

Unlike *SymRK*, *SHRK3* exhibits a presence-absence pattern unrelated to symbiotic associations, making it unlikely to be directly involved in AM (Delaux et al., 2014). This difference implies that *SymRK* and *SHRK3* are subject to stringent, yet distinct, functional constraints, despite their likely shared evolutionary origin. Notably, none of the tested species contained multiple copies of *SymRK* and *SHRK3* simultaneously (**Fig. 2**). Although this pattern should be interpreted cautiously given the current taxon sampling, it raises the possibility that simultaneous expansion of both clades may be constrained. Whether genetic interactions occur between these receptors remains to be determined (Iohannes and Jackson, 2023).

### Ectodomain protein characteristics differ among clades I–IV, suggesting distinct functional contributions

Among non-canonical protein variants, most variation was found within the ectodomains. This includes the loss of the MLD observed for SymRK in commelinids, as previously reported (**Fig. S5a, S9**) (Markmann et al., 2008; Miyata et al., 2023), as well as a variation in LRR number that contributes to differences in protein length (**Fig. S5b**). The observed contrast between ectodomain and intracellular domain diversity is consistent with previously reported asymmetry in LRR-RLK evolution (Man et al., 2023). This conceptually reflects the flexibility of the extracellular sensory domain in detecting diversified signals, while the intracellular domain maintains a conserved signal transduction function.

In SymRK, the release of the MLD generates a truncated receptor proteoform with increased affinity for Nod Factor Receptor 5, thereby promoting rhizobial symbiosis (Antolín-Llovera et al., 2014). The GDPC motif is both required for MLD release and a fingerprint of this receptor family. Analysis of the motif and its flanking regions has revealed clade-specific signatures (**Fig. S9c**). These signatures may differentially affect the cleavage, as was recently reported for SymRK (Chakrabarti et al., 2026). By contrast, in the *A. thaliana* clade IV member IOS1, the MLD has been suggested to contribute to receptor retention in the endoplasmic reticulum and suppression of the pathogen-induced unfolded protein response (Giordano et al., 2022).

Notably, the MLD also shares homology with *Cr*RLK1L proteins, which are involved in multiple aspects of plant growth, development, reproduction, and defence (Yang et al., 2021). In *Cr*RLK1Ls, these functions are mediated through the perception of RALF (Rapid Alkalinization Factor) peptides and interactions with cell wall components, either directly or via cell wall-associated proteins (Schade et al., 2025). It remains to be determined how the observed MLD diversity across clades I–IV relates to receptor function in plants, and whether MLD-dependent mechanisms are conserved among MLD-LRR-RLKs or distinct from those described for *Cr*RLK1Ls.

### Species-specific expansions are associated with gene tandem duplications in clade IV

The size of the SymRK receptor family varies among species, primarily due to lineage-specific heterogeneity in clades IV-X as reported previously for LRR-RLKs (Fischer et al., 2016; Kileeg et al., 2023; Shiu and Bleecker, 2003). Clades V-X represent lineage-specific expansion restricted to the non-flowering species in our sample. In some cases, such as the aquatic ferns *Azolla filiculoides* and *Ceratopteris brichardii*, these expansions could be related to particular adaptations to ecological niches or developmental contexts. Compared to vascular plants, bryophytes exhibit distinct patterns of diversification for accessory gene families, including *de novo* gene birth and horizontal gene transfer (Dong et al., 2025). Such differences in genome evolution may help explain the observed lower proportion of tandemly arrayed MLD-LRR-RLKs in bryophytes compared to vascular plants.

Closely related angiosperm species differ in the number of clade IV members, driven by uneven gene duplication events through mechanisms such as unequal crossover (Panchy et al., 2016). Gene duplication can lead to neofunctionalization and subfunctionalization due to relaxed selective pressure on the duplicated copy, providing a pool of targets for selection and adaptive evolution (Hanada et al., 2008; Panchy et al., 2016). Clade IV expansion is positively correlated with the number of genes located within tandem arrays. Tandem arrays represent genomic regions that generate receptor repertoire diversity and serve as a source of innovation for plant environmental adaptations. In *A. thaliana,* cluster D on chromosome five is part of a quantitative trait locus involved in root responses to *Pseudomonas* isolate 569, influencing root system architecture (Copeland et al., 2025, preprint). Moreover, the *NS1* and *NS2* loci in *M. truncatula* are part of a variable array of *MLD-LRR-RLKs,* which confer host specificity in the interaction with rhizobia (Liu et al., 2022; Yu et al., 2024).

Expansion via tandem arrays was especially pronounced in *A. thaliana,* where ∼70% of clade IV genes are clustered in tandem arrays. This reflects a broader trend of increased representation of MLD-LRR-RLKs observed in Brassicales (Kileeg et al., 2023). The dynamic nature of these clusters is also evident at the population level in *A. thaliana*. While nearly all single-dispersed *MLD-LRR-RLKs* are part of the conserved core or almost core categories and thus conserved across accessions, around 50% of *MLD-LRR-RLKs* exhibit presence-absence variation across accessions.

Interestingly, pangenome and phylogenetic analysis of clade A members in *A. thaliana* revealed that individual genes within the cluster expand through radiation. Similar patterns have previously been described in *A. thaliana* for nucleotide-binding domain and leucine-rich repeat receptors (NLRs) (Lee and Chae, 2020). Cluster radiation has been linked to instances of hybrid incompatibility. For example, the hybrids between the *A. thaliana* accession Bla-1 and Sha exhibit incompatibility and developmental abnormalities due to the genetic interaction between different *OAK* alleles (Smith et al., 2011). Therefore, the observed asymmetry in gene expansions may result from certain gene combinations that create stable phenotypes, while the incompatible combinations are naturally out-selected.

### Signatures of transcriptional diversification detected within clustered MLD-LRR-RLKs in *A. thaliana*

Analysis of transcriptional patterns revealed that, in *A. thaliana*, previously characterised *MLD-LRR-RLKs* (e.g. *FRK1*, *IOS1*) are comparatively highly expressed in shoots, whereas most clade IV genes show preferential expression in roots or seedlings. The biological significance of this preferential expression pattern remains to be determined. Nevertheless, the expression profile combined with responsiveness to biotic conditions, including root pathogens and commensals (**Fig. 5**), suggests that some members of this receptor family are transcriptionally integrated into biotic-response programmes.

Diversified expression profiles in tandem array clusters suggest that these genes may not be transcriptionally redundant. This could be due to duplicate genes missing portions of the original regulatory regions or evolving new regulatory regions, a hallmark of transcriptional sub-functionalisation. Partial exceptions include a subset of cluster A genes, which are transcriptionally co-regulated in shoots.

The trends reported here should be viewed as an initial framework to guide future functional analyses of this receptor family. Building on recent advances in single-cell analyses of immune responses (Nobori et al., 2025; Wang et al., 2026), future studies should focus on profiling the expression of these genes at cell-type resolution and under defined biotic conditions to better resolve their spatial and stimulus-specific expression patterns.

## Conclusions

SymRK is a conserved representative of the MLD-LRR-RLK receptor subfamily and served here as the founding reference member to define the SymRK receptor family. We found contrasting evolutionary patterns among MLD-LRR-RLK subfamily members, which grouped into four major and six additional lineage-specific clades. Clades I–III, including SymRK and its closest paralogs, are highly conserved and show limited gene copy number variation, consistent with stronger evolutionary constraint. In contrast, clade IV is more variable and characterised by lineage-specific differences in gene copy numbers, largely due to tandem duplications within gene clusters. These patterns are further evident at the population level in *A. thaliana,* highlighting the contribution of tandem arrays to receptor family dynamics. Finally, analysis of transcriptional expression patterns revealed that genes within the same cluster do not necessarily respond in a coordinated manner. *MLD-LRR-RLKs* show general transcriptional responsiveness to biotic conditions, with a subset of genes being repressed and the strongest responses observed in the shoot for the core genes of clade IV. Together, our results reveal contrasting evolutionary trajectories within the same receptor family and illustrate how plants may combine conservation of selected receptor lineages with diversification of others, providing a framework for future functional studies of MLD-LRR-RLK.

## Methods

### Phylogenetic analysis

#### Genome selection and sequence preparation

Genomes were downloaded from sources listed in their corresponding publications, or directly from NCBI and Phytozome (Suppl. Table 1). Where primary transcript files were supplied as part of the source genome annotation files, these were used for downstream analysis, otherwise the longest isoforms were filtered and extracted from the genome annotation files using the AGAT package (Dainat et al., 2026).

Orthogroups were assigned using OrthoFinder v2 (Emms and Kelly, 2019). All genes from orthogroups containing annotated SymRK/SHRK homologs in *L. japonicus* and *A. thaliana* were included for further filtering steps, independently of domain composition or length.

As an orthogonal approach to identify all MLD-LRR-RLKs, we performed a protein domain-based search of the original proteomes using HMMER 3.4 (Eddy, 2011)(see below). The identified proteins were combined with those identified by OrthoFinder to make up the full set of MLD-LRR-RLKs.

#### Tree-building

To build trees of all sequences, sequences were aligned using MAFFT (Katoh et al., 2002). Alignments were trimmed using BMGE (Criscuolo and Gribaldo, 2010) and trees were built using raxml-ng using the JTT+G model. For the unfiltered tree, sequences shorter than 1200 residues were aligned first, and longer sequences were added using the initial alignment as an anchor. Following the clade definition, sequences from each clade were realigned using MAFFT and built using IQ-Tree with 1000 bootstraps, using the standard ModelFinder and the —rate, —alninfo and —asr options. Trees were visualised using iTOL ver. 7.4 (Letunic and Bork, 2024).

#### Protein domain identification

To identify MLD-LRR-RLKs on the protein sequences, we used HMMSearch from HMMER 3.4 with the following profiles for each domain: PF12819 (Malectin-like), PF00069 (PKinase) and PF07714 (PK_Tyr_Ser-Thr). We filtered for an e-value of less than 0.001 and with an aligned region of at least half the length of the domain profile. In the case of the Kinase domains (PF00069, PF07714), we kept the best hit to avoid redundancy. Signal peptide and transmembrane regions were predicted using DeepTMHMM and used to determine the intra– and extracellular domains. We predicted LRRs using PhytoLRR (Chen, 2021). and selected candidate LRRs with a score higher than 7 and presenting at least 2 adjacent LRRs. Lastly, we filtered for proteins with the expected protein domain order, MLD, LRR, TM and Kinase domain. Canonical MLD-LRR-RLKs were defined as genes with high-confidence hits across all domains: MLD, LRR, and Kinase. Any homolog with any domain missing, or other predicted domains present, was labelled as non-canonical.

#### Protein domain analysis

Domain-wise percent similarity was calculated using the —percent-id option in clustal omega to generate a distance matrix for each group of sequences. The mean one-against-all percent identity was calculated as the row-wise mean of each row in the matrix. The same approach was used for all species as for within-angiosperm and within-non-angiosperm comparisons.

Sequence logos of the xDPC motif were generated using sequence alignments starting at – 1 from the conserved W residue. Logos were generated using WebLogo (Crooks et al., 2004).

#### Synteny analysis

Local synteny analysis for 69 *Arabidopsis thaliana* accessions was performed as described before (Lian et al., 2024). Three genomic regions encoding clusters of MLD-LRR-RLKs have been analysed, including regions 20 kb upstream and downstream of the cluster.

#### Gene cluster analysis

All MLD-LRR-RLKs included in the phylogenetic analysis were considered. The analysis focused on clades IV-X, which have expanded sets of homologous genes. For each clade, we identified the genomic positions of the corresponding genes and searched for sets of genes separated by no more than two intervening genes. The sets of genes were classified in three categories: (i) *tandem paralogs*, referring to two homologous genes located directly adjacent to each other; (ii) *proximal paralogs*, defined as duplicated genes separated by one or two intervening non-homologous genes; and (iii) *tandem clusters*, defined as groups of more than two genes formed a group with no more than two non-homologous genes between the members.

#### Gene expression analysis

General expression profiles for different tissues and biotic interactions were downloaded from: https://plantrnadb.com/athrdb/ (Yu et al., 2022). The tables contain the FPKM values for each gene in each library. Additionally, we obtained a table containing metadata, including project ID, genotype, ecotype, and tissue of origin. The tissue types were taken from (Yu et al., 2022). Using this information, we filtered for libraries and projects limited to wild-type *A. thaliana* accession Col-0. Libraries with low numbers of mapping reads were discarded. For the tissue-specific expression, this resulted in 878 libraries. The FPKM values were transformed to log2(FPKM+1) and batch-corrected using limma (v 3.64.3). After batch correction, we calculated the median expression level for each tissue to limit outlier bias.

We also obtained tissue-level gene expression (TMM counts) from www.travadb.org, based on (Klepikova et al., 2016).

Additionally, using metadata and annotations from (Yu et al., 2022), we selected libraries focused on interactions with diverse organisms. Transcriptomic projects without a control sample or that used a different accession background than Col-0 were discarded. The final dataset contains 372 transcriptomic profiles of *A. thaliana* with the following organisms: *Hyaloperonospora arabidopsidis, Botrytis cinerea, Colletotrichum tofieldiae, Heterodera schachti, Pseudomonas syringae, Fusarium graminearum, Rhizoctonia solani, Heterodera schachtii, Plutella xylostella, Pseudomonas simiae, Bacterial strain SA188, Microbacterium sp*.

The datasets were divided into three main groups based on their tissue of origin and to account for tissue-specific transcriptional responses: shoots, roots and seedlings.

For each project and tissue, and to account for variation across the different experimental designs, we calculated the log2 fold change of the medians between the treated and control samples by weighting by precision using the following formulas.

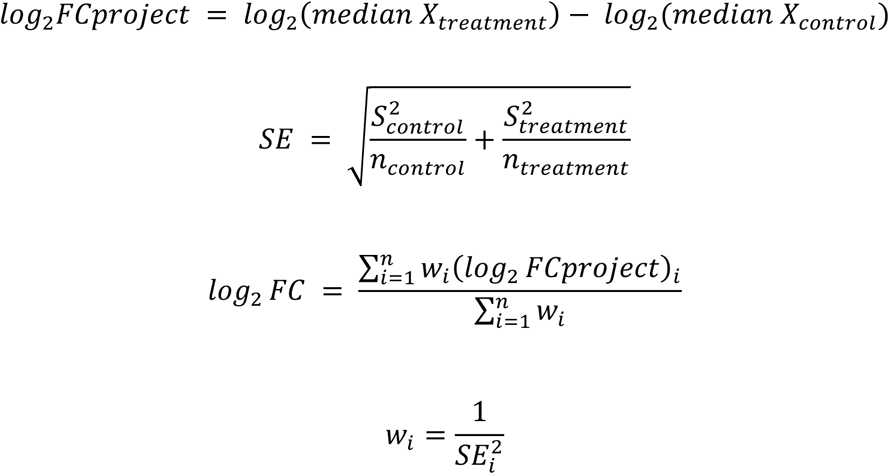

#### Presence-absence variation analysis of MLD-LRR-RLKs

Using all proteomes from the 69 accessions, we inferred orthogroups with OrthoFinder v2 (Emms and Kelly, 2019). In parallel, we identified all MLD-LRR-RLKs in 69 *A. thaliana* accessions as described previously. We further selected the orthogroups containing the canonical MLD-LRR-RLKs.

Most of the MLD-LRR-RLKs orthogroups were formed from single-copy orthologs. When an accession had multiple orthologs within a single orthogroup, we split those copies into new orthogroups. We repeated this process until we formed orthogroups containing only single genes per accession. From these orthogroups, we quantified the ortholog frequency across the accessions and assigned the following categories depending on the number of accessions per orthogroup: core if the ortholog is present in all accession, almost core if found in 62-68 accessions (90%), shell if 13-61 accessions (20-89%) carried the ortholog, cloud if the ortholog is present in 2-12 accessions (∼2 – 19%) and private if a gene is present in only one ecotype.

#### Data processing and visualization

Data processing and visualization was done using R (v4.5.1) ( Posit team, 2026).

## Declarations

### Authors’ contributions

This study was conceptualised by T.F-Ø, I.R., A.M., M.K.R-L., and K.P. M.P. initiated the research on SymRK and SHRKs. N.K. performed the preliminary analysis of the SymRK and SHRK orthogroups. T.F-Ø., A.M and M.K.R-L. selected species for phylogenetic analysis. T.F-Ø. conducted the phylogenetic analysis of MLD-LRR-RLKs in land plants, defined their clades, performed domain annotations, analysed structural and sequence variability, prepared Figures 1-2 and Figures S1-S10. I.R. analysed mechanisms of gene duplication in clade IV, examined the conservation of MLD-LRR-RLKs across *A. thaliana* accessions, performed transcriptional analysis across tissues and under biotic stress, prepared Figures 3-5 and Figures S11, S15-S17. A.M. analysed transcriptional responses to elicitor treatment and prepared Figure S18. K.S. conceptualised the synteny analysis of clustered clade IV members, which was performed by Q.L., who also prepared Figures S12-14. T.F-Ø., I.R. and K.P. wrote the manuscript with major contributions from A.M. and M.K.R-L. M.P., M.K.R-L., and K.P. acquired funding for this study.

## List of abbreviations

RLK: Receptor-like Kinase
MLD-LRR-RLK: Malectin-like Domain Leucine-rich Repeat Receptor-like Kinase
SymRK: Symbiosis Receptor-like Kinase
SHRK: SymRK-homologous Receptor-like Kinase
ECD: Extracellular Domain
ICD: Intracellular Domain
LRR-RLK: Leucine-rich Repeat Receptor-like Kinase
AM: Arbuscular Mycorrhiza
RNS: Root Nodule Symbiosis
*Hpa*: *Hyaloperonospora arabidopsidis*
SIMP1: Salt-Induced Malectin-like domain-containing Protein 1
IOS1: Impaired Oomycete Susceptibility 1
OAK1: Outgrowth-Associated protein Kinase 1
SIF: Stress Induced Factor
RHS: Root Hair Specific
BAK1: Brassinosteroid Insensitive 1 Associated Kinase 1
PRR: Pattern Recognition Receptor

## Acknowledgements

The authors would like to thank Maxim Messerer for initial guidance on Orthofinder analysis. HPC computing was performed on the BioHPC hosted at Leibniz Rechenzentrum Munich funded by Deutsche Forschungsgemeinschaft (DFG) project number 450674345. In addition, the authors gratefully acknowledge the computational and data resources provided by the Leibniz Supercomputing Centre (www.lrz.de). The authors acknowledge that artificial intelligence tools were used exclusively for language and grammatical editing.

## Figures, tables and additional files

**Supplementary Figure 1:** Taxonomic tree of 46 plant species used for phylogenetic analysis of the SymRK receptor family.

**Supplementary Figure 2:** A pipeline for improving the quality of phylogenetic tree reconstruction in the SymRK receptor family.

**Supplementary Figure 3:** Bootstrap and Orthogroup support for SymRK receptor family clade definitions.

**Supplementary Figure 4:** *Lotus japonicus* encodes eight MLD-LRR-RLKs.

**Supplementary Figure 5:** SymRK homologs identified in selected species.

**Supplementary Figure 6:** Phylogenetic tree for clade II homologs identified in selected species.

**Supplementary Figure 7:** Phylogenetic tree for clade III homologs identified in selected species.

**Supplementary Figure 8:** Phylogenetic tree for clade IV homologs identified in selected species.

**Supplementary Figure 9:** Patterns of structural variation in members of the SymRK receptor family.

**Supplementary Figure 10:** Residue-level polymorphism reveals asymmetric evolutionary patterns between extracellular and intracellular domains of SymRK homologs.

**Supplementary Figure 11:** Species-specific gene duplication events drive the expansion of the MLD-LRR-RLK receptor family.

**Supplementary Figure 12:** Local synteny of clustered clade IV members on chromosome 1 in *Arabidopsis thaliana*.

**Supplementary Figure 13:** Local synteny of clustered clade IV members on chromosome 3 in *Arabidopsis thaliana*.

**Supplementary Figure 14:** Local synteny of clustered clade IV members on chromosome 5 in *Arabidopsis thaliana*.

**Supplementary Figure 15:** Most of *Arabidopsis thaliana MLD-LRR-RLKs* are expressed in the root.

**Supplementary Figure 16:** *MLD-LRR-RLKs* generally exhibit low expression levels across different tissues in *Arabidopsis thaliana*.

**Supplementary Figure 17:** A subset of *MLD-LRR-RLKs* exhibits minimal expression levels across different tissues in *Arabidopsis thaliana*.

**Supplementary Figure 18:** Transcriptional responses of *Arabidopsis thaliana* clade IV genes to elicitor treatments.

## References

1. Antolín-Llovera, M., Ried, Martina K. & Parniske, M. (2014). Cleavage of the SYMBIOSIS RECEPTOR-LIKE KINASE Ectodomain Promotes Complex Formation with Nod Factor Receptor 5. Current Biology, 24, 422–427.

2. Asai, T., Tena, G., Plotnikova, J., Willmann, M. R., Chiu, W.-L., Gomez-Gomez, L., Boller, T., Ausubel, F. M. & Sheen, J. (2002). MAP kinase signalling cascade in Arabidopsis innate immunity. Nature, 415, 977–983.

3. Badouin, H., Gouzy, J., Grassa, C. J., Murat, F., Staton, S. E., Cottret, L., Lelandais-Brière, C., Owens, G. L., Carrère, S., Mayjonade, B., Legrand, L., Gill, N., Kane, N. C., Bowers, J. E., Hubner, S., Bellec, A., Bérard, A., Bergès, H., Blanchet, N., Boniface, M.-C., Brunel, D., Catrice, O., Chaidir, N., Claudel, C., Donnadieu, C., Faraut, T., Fievet, G., Helmstetter, N., King, M., Knapp, S. J., Lai, Z., Le Paslier, M.-C., Lippi, Y., Lorenzon, L., Mandel, J. R., Marage, G., Marchand, G., Marquand, E., Bret-Mestries, E., Morien, E., Nambeesan, S., Nguyen, T., Pegot-Espagnet, P., Pouilly, N., Raftis, F., Sallet, E., Schiex, T., Thomas, J., Vandecasteele, C., Varès, D., Vear, F., Vautrin, S., Crespi, M., Mangin, B., Burke, J. M., Salse, J., Muños, S., Vincourt, P., Rieseberg, L. H. & Langlade, N. B. (2017). The sunflower genome provides insights into oil metabolism, flowering and Asterid evolution. Nature, 546, 148–152.

4. Bjornson, M., Pimprikar, P., Nürnberger, T. & Zipfel, C. (2021). The transcriptional landscape of *Arabidopsis thaliana* pattern-triggered immunity. Nat Plants, 7, 579–586.

5. Chakrabarti, D., Paul, A., Molla, F., Bhattacharyya, S., Das, S., Chakraborty, S., Ghosh, D., Biswas, A., Kundu, A., Sinharoy, S. & Dasgupta, M. (2026). Conserved hinge regions in SYMRK enable release of Malectin-like Domain for symbiont passage during rhizobia–legume symbiosis. The Plant Cell.

6. Chan, C., Panzeri, D., Okuma, E., Tõldsepp, K., Wang, Y.-Y., Louh, G.-Y., Chin, T.-C., Yeh, Y.-H., Yeh, H.-L., Yekondi, S., Huang, Y.-H., Huang, T.-Y., Chiou, T.-J., Murata, Y., Kollist, H. & Zimmerli, L. (2020). STRESS INDUCED FACTOR 2 Regulates Arabidopsis Stomatal Immunity through Phosphorylation of the Anion Channel SLAC1. The Plant Cell, 32, 2216–2236.

7. Chen, T. (2021). Identification and characterization of the LRR repeats in plant LRR-RLKs. BMC Mol Cell Biol, 22, 9.

8. Copeland, C., Logemann, E., Malisic, M., Amrhein, A., Valisi, A. & Schulze-Lefert, P. (2025). A multigenic quantitative trait locus underlies natural variation in *Arabidopsis thaliana* root system architecture and transcriptional responses to microbiota-derived *Pseudomonas*. bioRxiv, 2025.09.03.673899.

9. Criscuolo, A. & Gribaldo, S. (2010). BMGE (Block Mapping and Gathering with Entropy): a new software for selection of phylogenetic informative regions from multiple sequence alignments. BMC Evol Biol, 10, 210.

10. Crooks, G. E., Hon, G., Chandonia, J. M. & Brenner, S. E. (2004). WebLogo: a sequence logo generator. Genome Res, 14, 1188–90.

11. Cui, J., Hu, Y., Huang, Y., Guo, J., Xie, X., Zhang, H. & Cai, Y. (2022). Loss of Function Mutation of IOS1 in *Arabidopsis* Is More Sensitive to Salt Stress. Plant Molecular Biology Reporter, 40, 68–80.

12. Dainat, J., Cannoodt, R., Soares, A., García Ruano, D., Hereñú, D., Murray, D. K. D., Davis, E., Ugrin, I., Crouch, K., Soler, L., Pascal-Git, Zollman, Z. & Tayyrov. 2026. NBISweden/AGAT: AGAT v1.7.0 (v1.7.0). Zenodo.

13. De Smet, R., Adams, K. L., Vandepoele, K., Van Montagu, M. C. E., Maere, S. & Van De Peer, Y. (2013). Convergent gene loss following gene and genome duplications creates single-copy families in flowering plants. Proceedings of the National Academy of Sciences, 110, 2898–2903.

14. Defalco, T. A. & Zipfel, C. (2021). Molecular mechanisms of early plant pattern-triggered immune signaling. Molecular Cell, 81, 3449–3467.

15. Delaux, P.-M., Varala, K., Edger, P. P., Coruzzi, G. M., Pires, J. C. & Ané, J.-M. (2014). Comparative Phylogenomics Uncovers the Impact of Symbiotic Associations on Host Genome Evolution. PLOS Genetics, 10, e1004487.

16. Demchenko, K., Winzer, T., Stougaard, J., Parniske, M. & Pawlowski, K. (2004). Distinct roles of *Lotus japonicus* SYMRK and SYM15 in root colonization and arbuscule formation. New Phytologist, 163, 381–392.

17. Dievart, A., Gottin, C., Périn, C., Ranwez, V. & Chantret, N. (2020). Origin and Diversity of Plant Receptor-Like Kinases. Annu Rev Plant Biol, 71, 131–156.

18. Dong, S., Wang, S., Li, L., Yu, J., Zhang, Y., Xue, J.-Y., Chen, H., Ma, J., Zeng, Y., Cai, Y., Huang, W., Zhou, X., Wu, J., Li, J., Yao, Y., Hu, R., Zhao, T., Villarreal A, J. C., Dirick, L., Liu, L., Ignatov, M., Jin, M., Ruan, J., He, Y., Wang, H., Xu, B., Rozzi, R., Wegrzyn, J., Stevenson, D. W., Renzaglia, K. S., Chen, H., Zhang, L., Zhang, S., Mackenzie, R., Moreno, J. E., Melkonian, M., Wei, T., Gu, Y., Xu, X., Rensing, S. A., Huang, J., Long, M., Goffinet, B., Bowman, J. L., Van De Peer, Y., Liu, H. & Liu, Y. (2025). Bryophytes hold a larger gene family space than vascular plants. Nature Genetics, 57, 2562–2569.

19. Eddy, S. R. (2011). Accelerated Profile HMM Searches. PLOS Computational Biology, 7, e1002195.

20. Emms, D. M. & Kelly, S. (2019). OrthoFinder: phylogenetic orthology inference for comparative genomics. Genome Biology, 20, 238.

21. Endre, G., Kereszt, A., Kevei, Z., Mihacea, S., Kaló, P. & Kiss, G. B. (2002). A receptor kinase gene regulating symbiotic nodule development. Nature, 417, 962–6.

22. Escocard De Azevedo Manhães, A. M., Ortiz-Morea, F. A., He, P. & Shan, L. (2021). Plant plasma membrane-resident receptors: Surveillance for infections and coordination for growth and development. Journal of Integrative Plant Biology, 63, 79–101.

23. Feng, Y., Wu, P., Liu, C., Peng, L., Wang, T., Wang, C., Tan, Q., Li, B., Ou, Y., Zhu, H., Yuan, S., Huang, R., Stacey, G., Zhang, Z. & Cao, Y. (2021). Suppression of *Lj*BAK1-mediated immunity by SymRK promotes rhizobial infection in *Lotus japonicus*. Molecular Plant, 14, 1935–1950.

24. Fischer, I., Diévart, A., Droc, G., Dufayard, J.-F. & Chantret, N. (2016). Evolutionary Dynamics of the Leucine-Rich Repeat Receptor-Like Kinase (LRR-RLK) Subfamily in Angiosperms Plant Physiology, 170, 1595–1610.

25. Giordano, L., Schimmerling, M., Panabières, F., Allasia, V. & Keller, H. (2022). The exodomain of the impaired oomycete susceptibility 1 receptor mediates both endoplasmic reticulum stress responses and abscisic acid signalling during downy mildew infection of Arabidopsis. Molecular Plant Pathology, 23, 1783–1791.

26. Hajný, J., Prát, T., Rydza, N., Rodriguez, L., Tan, S., Verstraeten, I., Domjan, D., Mazur, E., Smakowska-Luzan, E., Smet, W., Mor, E., Nolf, J., Yang, B., Grunewald, W., Molnár, G., Belkhadir, Y., De Rybel, B. & Friml, J. (2020). Receptor kinase module targets PIN-dependent auxin transport during canalization. Science, 370, 550–557.

27. Hanada, K., Zou, C., Lehti-Shiu, M. D., Shinozaki, K. & Shiu, S.-H. (2008). Importance of Lineage-Specific Expansion of Plant Tandem Duplicates in the Adaptive Response to Environmental Stimuli Plant Physiology, 148, 993–1003.

28. He, J., Zhuang, Y., Li, C., Sun, X., Zhao, S., Ma, C., Lin, H. & Zhou, H. (2021). SIMP1 modulates salt tolerance by elevating ERAD efficiency through UMP1A-mediated proteasome maturation in plants. New Phytologist, 232, 625–641.

29. nn, U., Lau, K. & Hothorn, M. (2017). The Structural Basis of Ligand Perception and Signal Activation by Receptor Kinases. Annu Rev Plant Biol, 68, 109–137.

30. Hok, S., Allasia, V., Andrio, E., Naessens, E., Ribes, E., Panabières, F., Attard, A., Ris, N., Clément, M., Barlet, X., Marco, Y., Grill, E., Eichmann, R., Weis, C., Hückelhoven, R., Ammon, A., Ludwig-Müller, J., Voll, L. M. & Keller, H. (2014). The Receptor Kinase IMPAIRED OOMYCETE SUSCEPTIBILITY1 Attenuates Abscisic Acid Responses in Arabidopsis Plant Physiology, 166, 1506–1518.

31. Hok, S., Danchin, E. G. J., Allasia, V., Panabiéres, F., Attard, A. & Keller, H. (2011). An Arabidopsis (malectin-like) leucine-rich repeat receptor-like kinase contributes to downy mildew disease. Plant, Cell & Environment, 34, 1944–1957.

32. Hosseini, S., Schmidt, E. D. L. & Bakker, F. T. (2020). Leucine-rich repeat receptor-like kinase II phylogenetics reveals five main clades throughout the plant kingdom. Plant J, 103, 547–560.

33. Iohannes, S. D. & Jackson, D. (2023). Tackling redundancy: genetic mechanisms underlying paralog compensation in plants. New Phytologist, 240, 1381–1389.

34. Katoh, K., Misawa, K., Kuma, K. I. & Miyata, T. (2002). MAFFT: a novel method for rapid multiple sequence alignment based on fast Fourier transform. Nucleic Acids Research, 30, 3059–3066.

35. Kileeg, Z., Haldar, A., Khan, H., Qamar, A. & Mott, G. A. (2023). Differential expansion and retention patterns of LRR-RLK genes across plant evolution. Plant Direct, 7, e556.

36. Klepikova, A. V., Kasianov, A. S., Gerasimov, E. S., Logacheva, M. D. & Penin, A. A. (2016). A high resolution map of the *Arabidopsis thaliana* developmental transcriptome based on RNA-seq profiling. Plant J, 88, 1058–1070.

37. Lee, R. R. Q. & Chae, E. (2020). Variation Patterns of NLR Clusters in *Arabidopsis thaliana* Genomes. Plant Commun, 1, 100089.

38. Lehti-Shiu, M. D., Zou, C., Hanada, K. & Shiu, S.-H. (2009). Evolutionary History and Stress Regulation of Plant Receptor-Like Kinase/Pelle Genes Plant Physiology, 150, 12–26.

39. Letunic, I. & Bork, P. (2024). Interactive Tree of Life (iTOL) v6: recent updates to the phylogenetic tree display and annotation tool. Nucleic Acids Res, 52, W78–w82.

40. Lian, Q., Huettel, B., Walkemeier, B., Mayjonade, B., Lopez-Roques, C., Gil, L., Roux, F., Schneeberger, K. & Mercier, R. (2024). A pan-genome of 69 *Arabidopsis thaliana* accessions reveals a conserved genome structure throughout the global species range. Nature Genetics, 56, 982–991.

41. Liu, J., Wang, T., Qin, Q., Yu, X., Yang, S., Dinkins, R. D., Kuczmog, A., Putnoky, P., Muszyński, A., Griffitts, J. S., Kereszt, A. & Zhu, H. (2022). Paired *Medicago* receptors mediate broad-spectrum resistance to nodulation by *Sinorhizobium meliloti* carrying a species-specific gene. Proc Natl Acad Sci U S A, 119, e2214703119.

42. Man, J., Gallagher, J. P. & Bartlett, M. (2020). Structural evolution drives diversification of the large LRR-RLK gene family. New Phytologist, 226, 1492–1505.

43. Man, J., Harrington, T. A., Lally, K. & Bartlett, M. E. (2023). Asymmetric Evolution of Protein Domains in the Leucine-Rich Repeat Receptor-Like Kinase Family of Plant Signaling Proteins. Molecular Biology and Evolution, 40.

44. Markmann, K., Giczey, G. & Parniske, M. (2008). Functional Adaptation of a Plant Receptor-Kinase Paved the Way for the Evolution of Intracellular Root Symbioses with Bacteria. PLOS Biology, 6, e68.

45. Miyata, K., Hosotani, M., Akamatsu, A., Takeda, N., Jiang, W., Sugiyama, T., Takaoka, R., Matsumoto, K., Abe, S., Shibuya, N. & Kaku, H. (2023). *Os*SYMRK Plays an Essential Role in AM Symbiosis in Rice (*Oryza sativa*). Plant and Cell Physiology, 64, 378–391.

46. Ngou, B. P. M., Wyler, M., Schmid, M. W., Kadota, Y. & Shirasu, K. (2024). Evolutionary trajectory of pattern recognition receptors in plants. Nature Communications, 15, 308.

47. Nobori, T., Monell, A., Lee, T. A., Sakata, Y., Shirahama, S., Zhou, J., Nery, J. R., Mine, A. & Ecker, J. R. (2025). A rare PRIMER cell state in plant immunity. Nature, 638, 197–205.

48. Panchy, N., Lehti-Shiu, M. & Shiu, S. H. (2016). Evolution of Gene Duplication in Plants. Plant Physiol, 171, 2294–316.

49. Piet, Q., Droc, G., Marande, W., Sarah, G., Bocs, S., Klopp, C., Bourge, M., Siljak-Yakovlev, S., Bouchez, O., Lopez-Roques, C., Lepers-Andrzejewski, S., Bourgois, L., Zucca, J., Dron, M., Besse, P., Grisoni, M., Jourda, C. & Charron, C. (2022). A chromosome-level, haplotype-phased *Vanilla planifolia* genome highlights the challenge of partial endoreplication for accurate whole-genome assembly. Plant Communications, 3, 100330.

50. Radhakrishnan, G. V., Keller, J., Rich, M. K., Vernié, T., Mbadinga Mbadinga, D. L., Vigneron, N., Cottret, L., Clemente, H. S., Libourel, C., Cheema, J., Linde, A.-M., Eklund, D. M., Cheng, S., Wong, G. K. S., Lagercrantz, U., Li, F.-W., Oldroyd, G. E. D. & Delaux, P.-M. (2020). An ancestral signalling pathway is conserved in intracellular symbioses-forming plant lineages. Nature Plants, 6, 280–289.

51. Ried, M. K., Banhara, A., Hwu, F. Y., Binder, A., Gust, A. A., Höfle, C., Hückelhoven, R., Nürnberger, T. & Parniske, M. (2019). A set of Arabidopsis genes involved in the accommodation of the downy mildew pathogen *Hyaloperonospora arabidopsidis*. PLoS Pathog, 15, e1007747.

52. Sageman-Furnas, K., Nurmi, M., Contag, M., Plötner, B., Alseekh, S., Wiszniewski, A., Fernie, A. R., Smith, L. M. & Laitinen, R. a. E. (2022). *A. thaliana* Hybrids Develop Growth Abnormalities through Integration of Stress, Hormone and Growth Signaling. Plant and Cell Physiology, 63, 944–954.

53. Schade, S., Leicher, H., Von Arx, M., Monte, I., Gronnier, J. & Stegmann, M. (2025). The interplay of RALF structural and signaling functions in plant-microbe interactions. PLOS Pathogens, 21, e1013588.

54. Shiu, S.-H. & Bleecker, A. B. (2001a). Plant Receptor-Like Kinase Gene Family: Diversity, Function, and Signaling. Science’s STKE, 2001, re22–re22.

55. Shiu, S.-H. & Bleecker, A. B. (2001b). Receptor-like kinases from Arabidopsis form a monophyletic gene family related to animal receptor kinases. Proceedings of the National Academy of Sciences, 98, 10763–10768.

56. Shiu, S.-H. & Bleecker, A. B. (2003). Expansion of the Receptor-Like Kinase/Pelle Gene Family and Receptor-Like Proteins in Arabidopsis Plant Physiology, 132, 530–543.

57. Smith, L. M., Bomblies, K. & Weigel, D. (2011). Complex Evolutionary Events at a Tandem Cluster of *Arabidopsis thaliana* Genes Resulting in a Single-Locus Genetic Incompatibility. PLOS Genetics, 7, e1002164.

58. Stracke, S., Kistner, C., Yoshida, S., Mulder, L., Sato, S., Kaneko, T., Tabata, S., Sandal, N., Stougaard, J., Szczyglowski, K. & Parniske, M. (2002). A plant receptor-like kinase required for both bacterial and fungal symbiosis. Nature, 417, 959–62.

59. Team, P. 2026. RStudio: Integrated Development Environment for R.: Posit Software, PBC, Boston, MA.

60. Vaattovaara, A., Brandt, B., Rajaraman, S., Safronov, O., Veidenberg, A., Luklová, M., Kangasjärvi, J., Löytynoja, A., Hothorn, M., Salojärvi, J. & Wrzaczek, M. (2019). Mechanistic insights into the evolution of DUF26-containing proteins in land plants. Commun Biol, 2, 56.

61. Wang, S., Bezrukov, I., Wu, P.-J., Gauß, H., Timmermans, M. & Weigel, D. (2026). Cell-type-specific gating of gene regulatory modules as a hallmark of early immune responses in Arabidopsis leaves. New Phytologist, 250, 2007–2025.

62. Won, S.-K., Lee, Y.-J., Lee, H.-Y., Heo, Y.-K., Cho, M. & Cho, H.-T. (2009). cis-Element– and Transcriptome-Based Screening of Root Hair-Specific Genes and Their Functional Characterization in Arabidopsis Plant Physiology, 150, 1459–1473.

63. Xu, P., Xu, S. L., Li, Z. J., Tang, W., Burlingame, A. L. & Wang, Z. Y. (2014). A brassinosteroid-signaling kinase interacts with multiple receptor-like kinases in Arabidopsis. Mol Plant, 7, 441–4.

64. Yang, H., Wang, D., Guo, L., Pan, H., Yvon, R., Garman, S., Wu, H.-M. & Cheung, A. Y. (2021). Malectin/Malectin-like domain-containing proteins: A repertoire of cell surface molecules with broad functional potential. The Cell Surface, 7, 100056.

65. Yeh, Y.-H., Panzeri, D., Kadota, Y., Huang, Y.-C., Huang, P.-Y., Tao, C.-N., Roux, M., Chien, H.-C., Chin, T.-C., Chu, P.-W., Zipfel, C. & Zimmerli, L. (2016). The Arabidopsis Malectin-Like/LRR-RLK IOS1 Is Critical for BAK1-Dependent and BAK1-Independent Pattern-Triggered Immunity. The Plant Cell, 28, 1701–1721.

66. Yu, X., Liu, J., Qin, Q., Zribi, I., Yu, J., Yang, S., Dinkins, R. D., Fei, Z., Kereszt, A. & Zhu, H. (2024). Species-specific microsymbiont discrimination mediated by a *Medicago* receptor kinase. Science Advances, 10, eadp6436.

67. Yu, Y., Zhang, H., Long, Y., Shu, Y. & Zhai, J. (2022). Plant Public RNA-seq Database: a comprehensive online database for expression analysis of ∼45 000 plant public RNA-Seq libraries. Plant Biotechnology Journal, 20, 806–808.

68. Yuan, N., Yuan, S., Li, Z., Zhou, M., Wu, P., Hu, Q., Mendu, V., Wang, L. & Luo, H. (2018). STRESS INDUCED FACTOR 2, a Leucine-Rich Repeat Kinase Regulates Basal Plant Pathogen Defense Plant Physiology, 176, 3062–3080.

