## Supplemental Data 1 for "Evolutionary diversification of the SymRK receptor family in land plants"

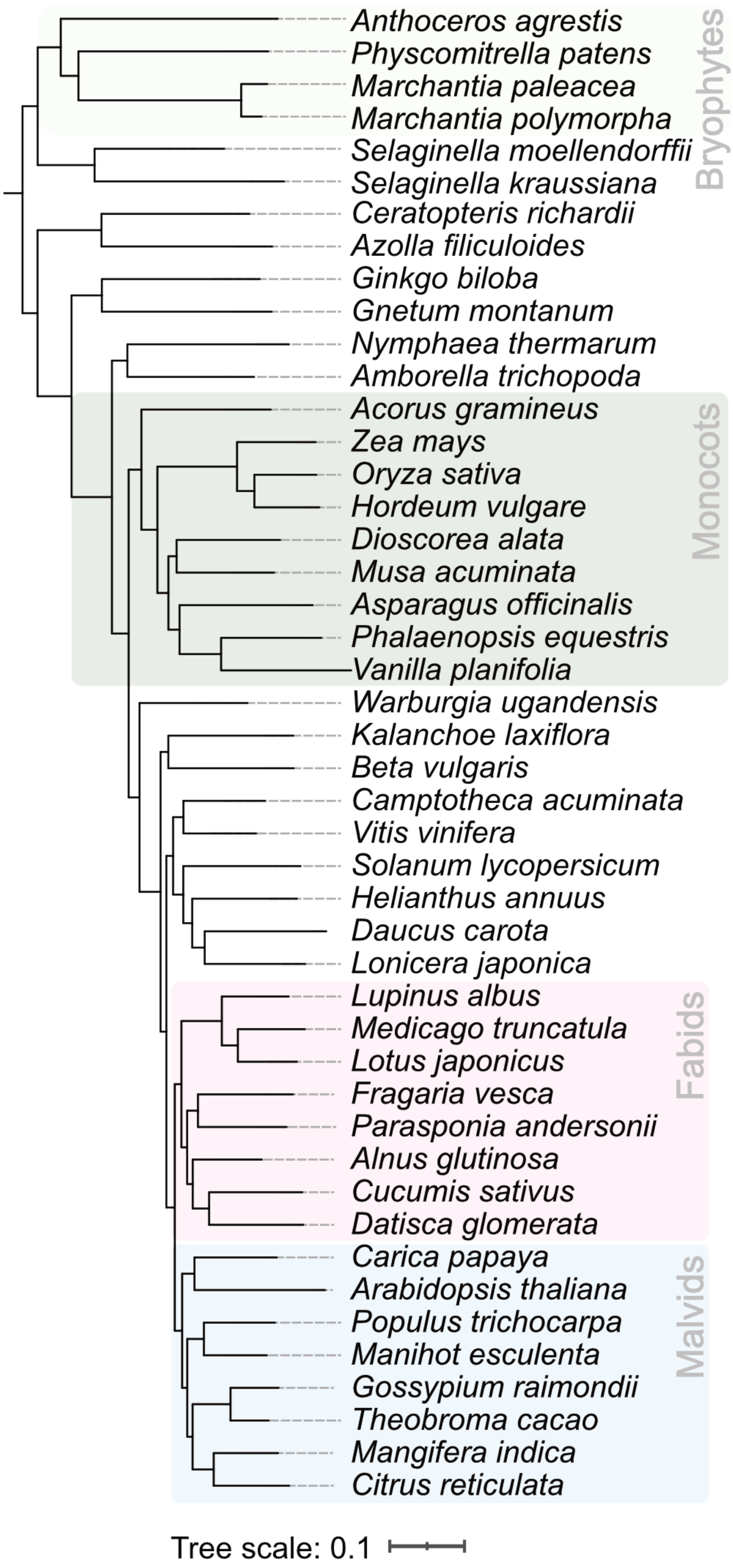


**Supplementary Figure 1: Taxonomic tree of 46 plant species used for phylogenetic analysis of the SymRK receptor family.** Major plant lineages (Bryophytes, Monocots, Fabids and Malvids) are highlighted. The tree was built using OrthoFinder. See Table S1 for details on the genome version used for the analysis.

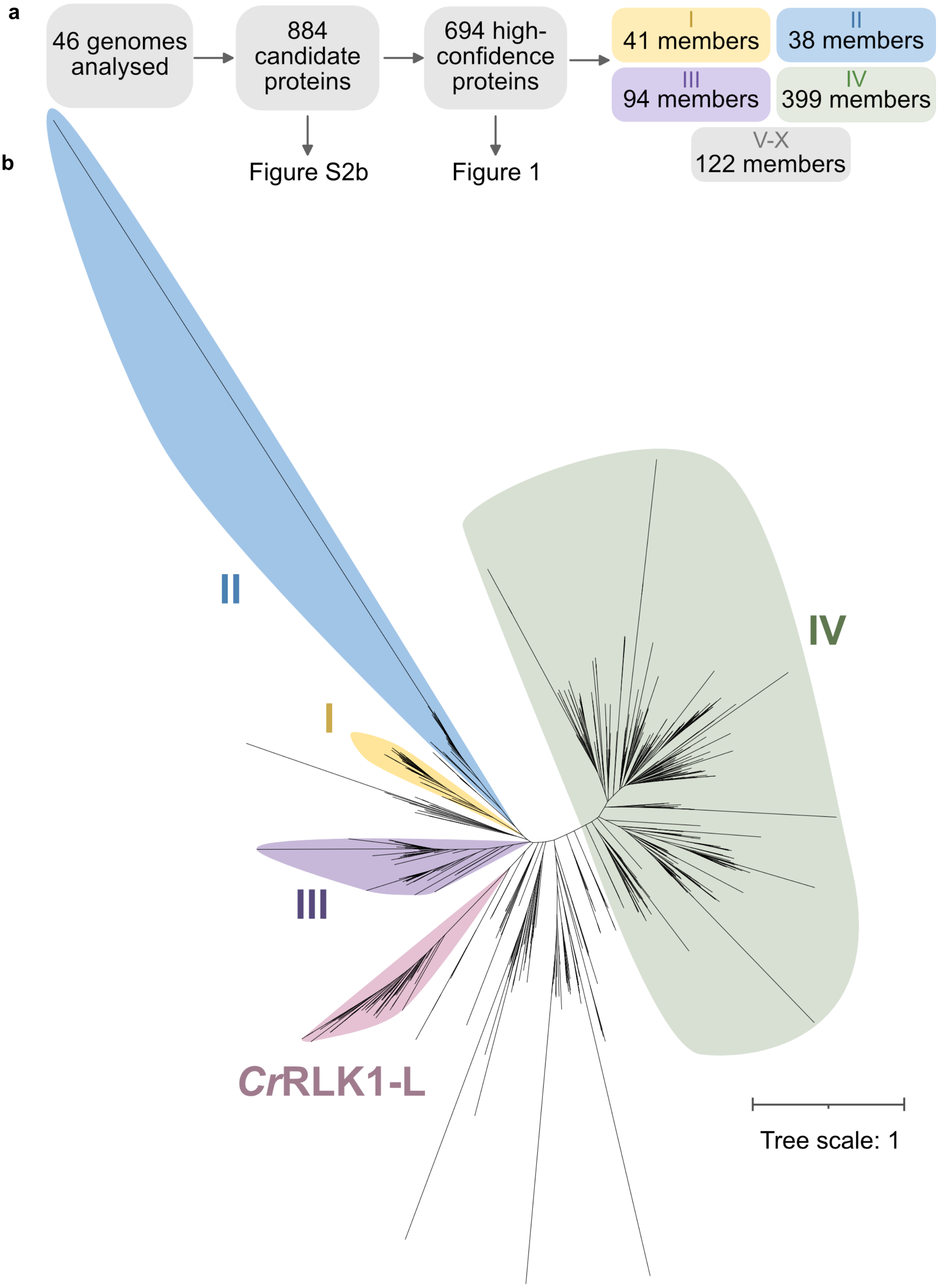


**Supplementary Figure 2: A pipeline for improving the quality of phylogenetic tree reconstruction in the SymRK receptor family. a -** A total of 884 putative MLD-LRR-RLK protein sequences with homology to SymRK were retrieved in 46 plant genomes using OrthoFinder and HMM-based domain searches, of which 694 were identified as originating from high-confidence genes after filtering steps (see Materials and Methods for details). These sequences were used to infer the phylogenetic tree in Figure 1 and to assign major clades. **b –** Phylogenetic tree of the SymRK receptor family built using all 884 candidate protein sequences prior to any filtering steps. The annotated clades reflect the closest matches to those annotated in Figure 1. Sequences were aligned using MAFFT, trimmed using BMGE and the tree was built using RAXML-NG.

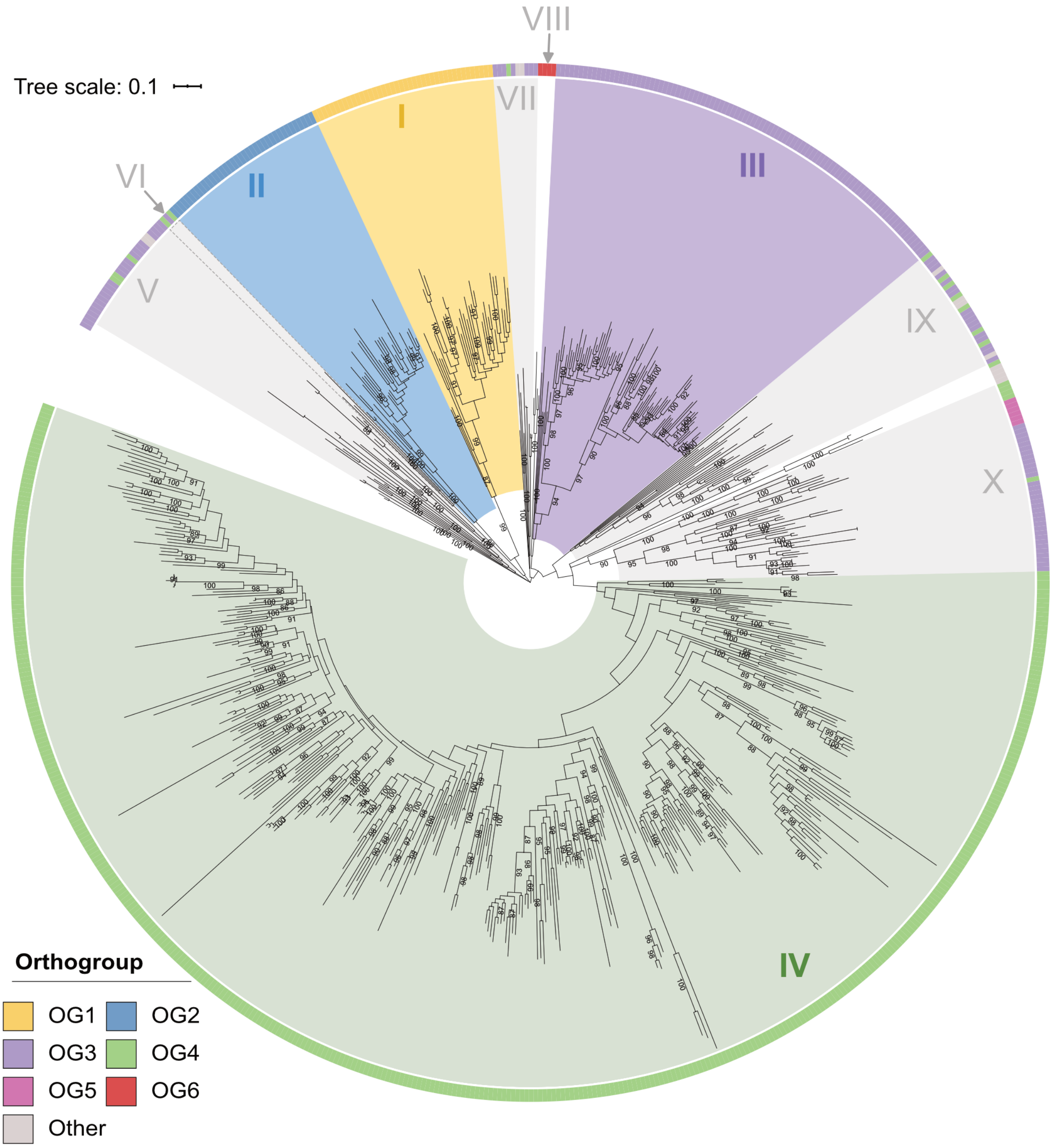


**Supplementary Figure 3: Bootstrap and Orthogroup support for SymRK receptor family clade definitions.** The same maximum-likelihood phylogenetic tree of proteins homologous to SymRK as shown in Figure 1, with the main clades annotated together with their bootstrap support. Clade definitions are based on bootstrap values and monophyletic orthogroup distribution. Bootstrap values above 85 are displayed. The outer ring colour corresponds to orthogroup (OG) assignments. Angiosperm members of OG1-4 largely correspond to clades I-IV, respectively. Homologous receptors belonging to other orthogroups were identified in non-angiosperm species and found to be distributed between ten different orthogroups. Of these, only OG5 and OG6 contained more than two identified homologous receptors. The remaining orthogroups are shown in grey (other).

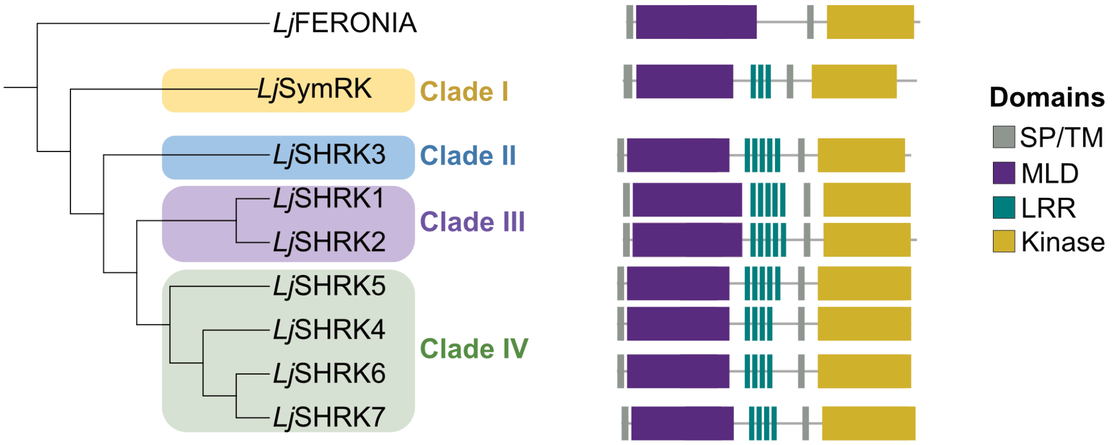


**Supplementary Figure 4: *Lotus japonicus* encodes eight MLD-LRR-RLKs**. These MLD-LRR-RLKs, referred to as SymRK and SHRK1-7, are distributed between the clades identified in Fig. 1 and are highlighted in the corresponding clade colours. Their predicted domain architectures are illustrated to the right. The *L. japonicus* FERONIA sequence was used as an outgroup for tree construction. SP – Signal Peptide, TM – Transmembrane domain, MLD – Malectin-like Domain, LRR – Leucine-rich Repeat, Kinase – Kinase domain.

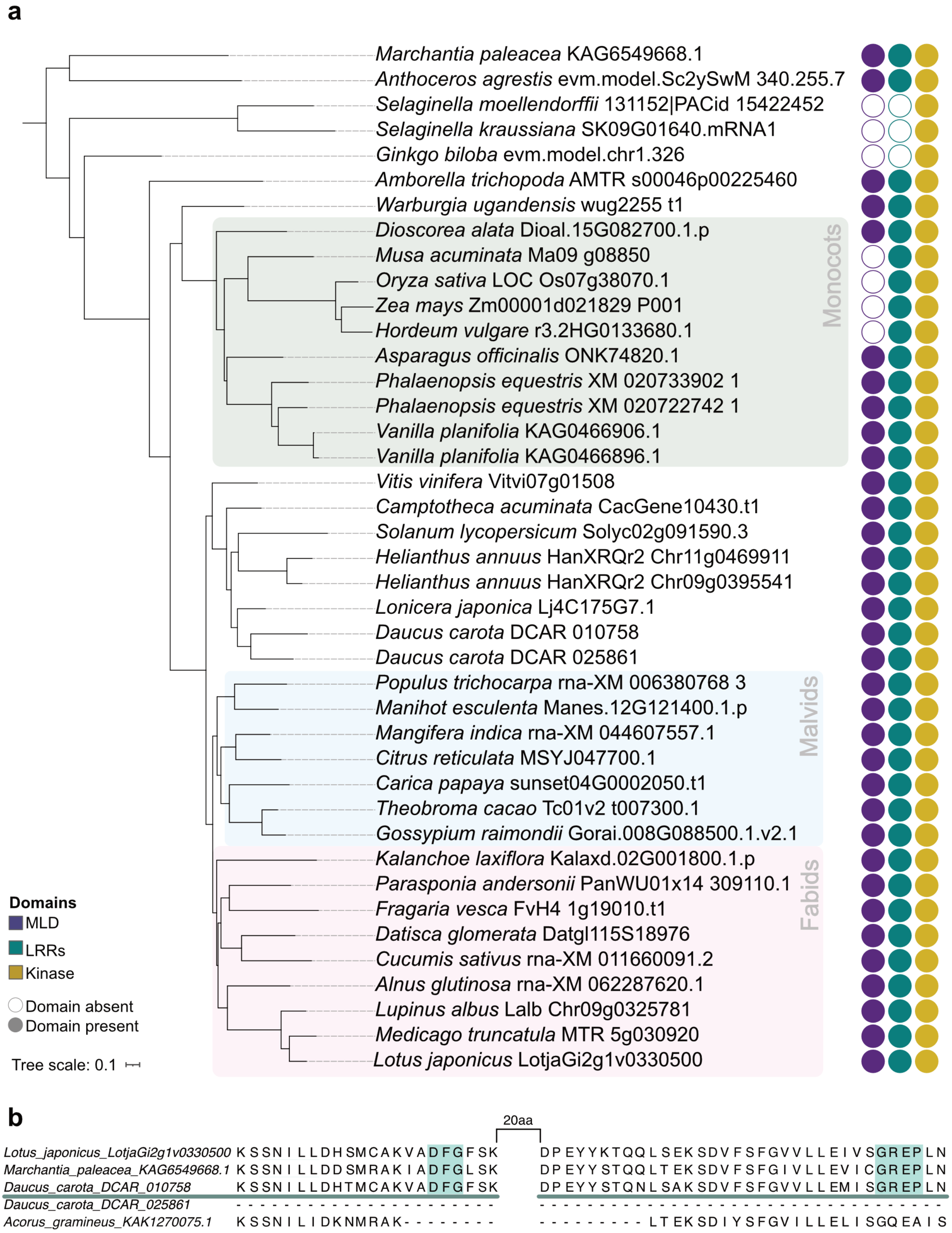


**Supplementary Figure 5: SymRK homologs identified in selected species. a** - A phylogenetic tree for SymRK homologs was constructed using IQtree based on protein sequences assigned to clade I. The predicted presence or absence of domains in each homolog is indicated on the right. MLD - Malectin-like Domain, LRRs - Leucine-rich Repeats and Kinase - Kinase domain. Major taxonomic groups are highlighted. **b** - *Daucus carota* and *Acorus gramineus* encode SymRK protein variants with deletions in the conserved kinase domain. Shown is an alignment over two short segments of SymRK kinase domain separated by 20 amino acids (aa) (amino acids 722-743 and 765-798 in *L. japonicus* SymRK) for selected homologs to highlight the deletions in key conserved motifs, DFG and GREP (highlighted in green). Note that *Daucus carota* encodes two SymRK variants, one full-length and one with a deletion in the kinase.

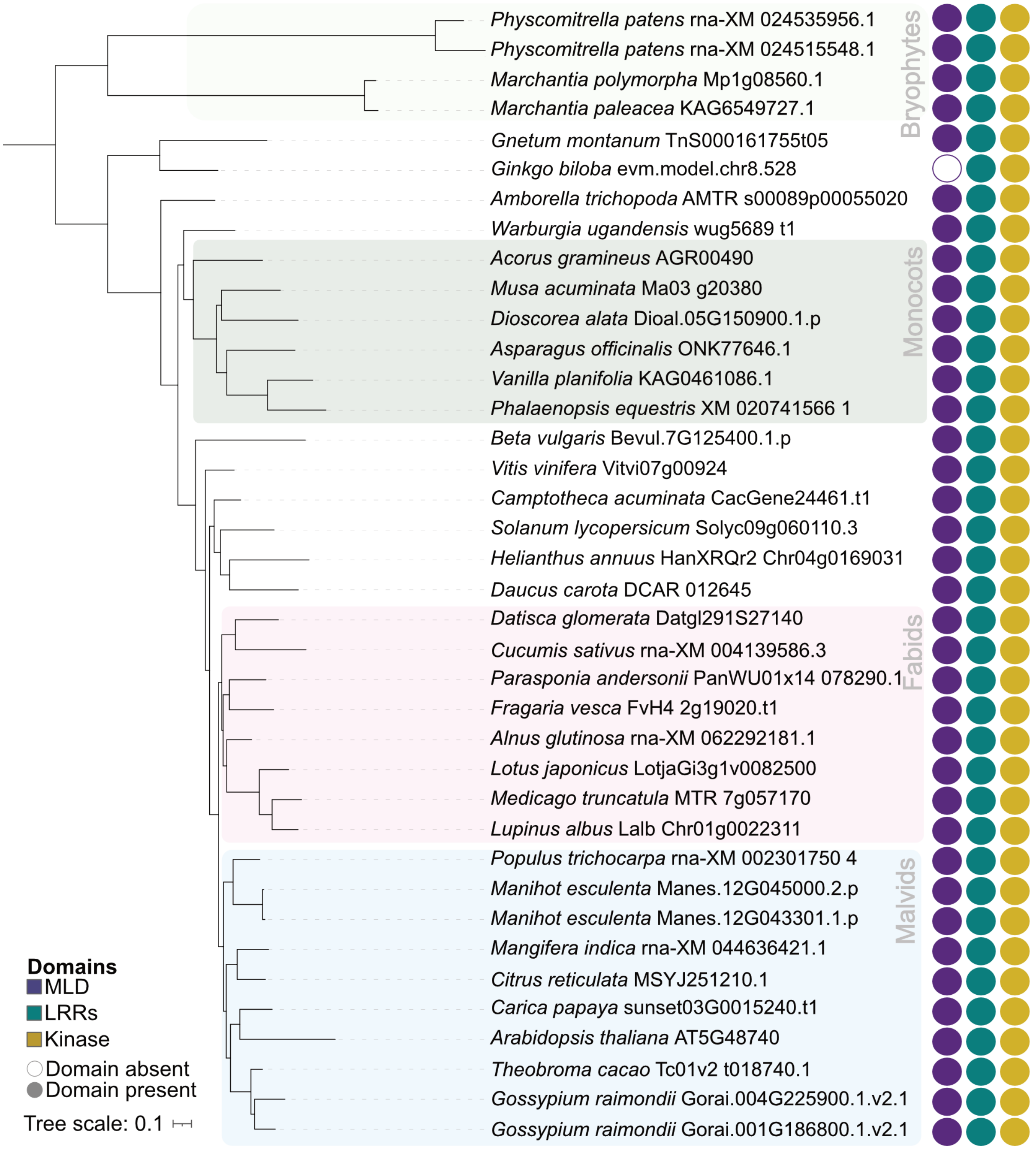


**Supplementary Figure 6: Phylogenetic tree for clade II homologs identified in selected species.** The tree was constructed using IQtree based on protein sequences assigned to clade II. The predicted presence or absence of domains in each homolog is indicated on the right. MLD - Malectin-like Domain, LRRs - Leucine-rich Repeats and Kinase - Kinase domain.

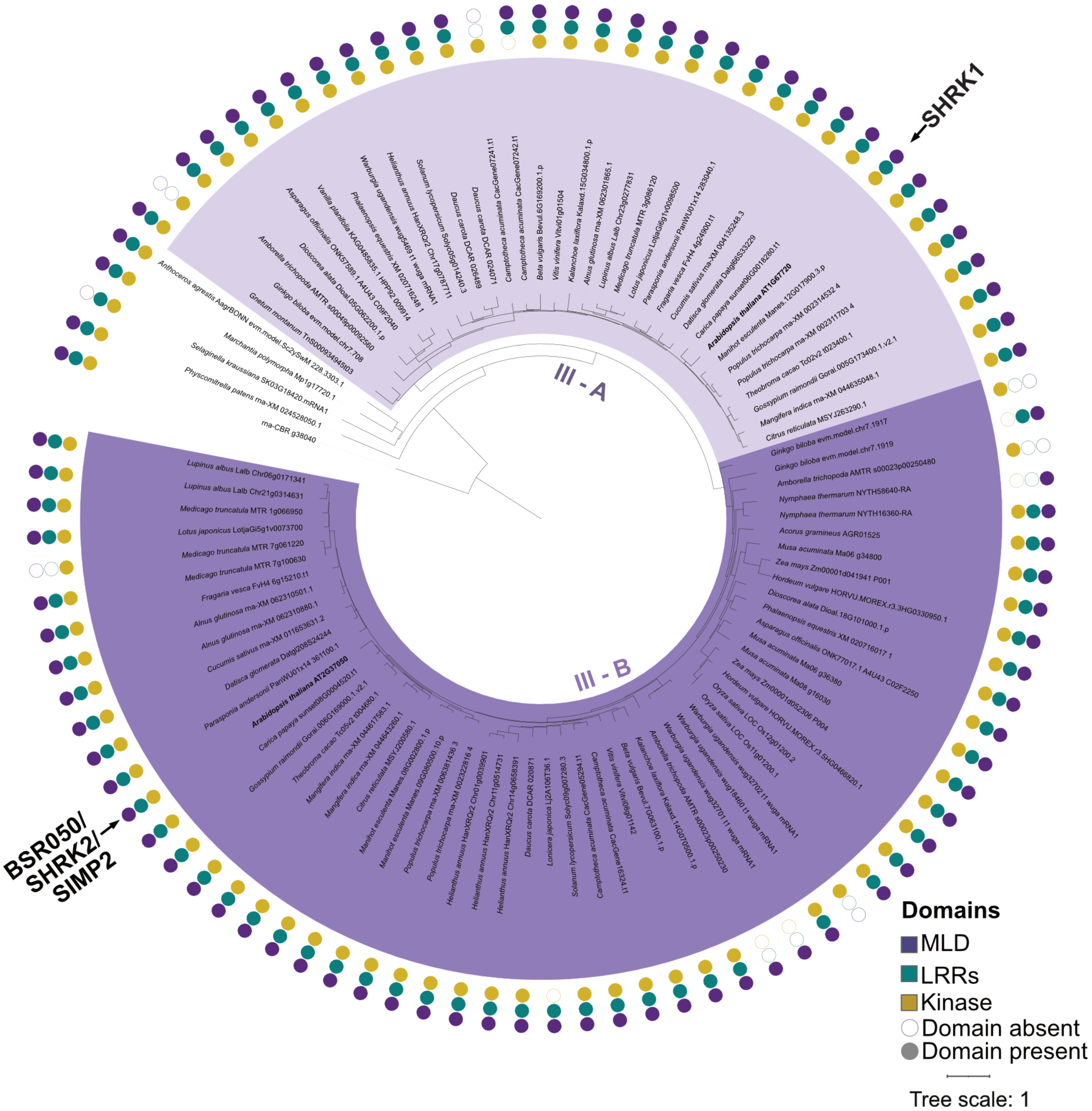


**Supplementary Figure 7: Phylogenetic tree for clade III homologs identified in selected species.** The tree was constructed using IQtree with protein sequences assigned to clade III. A FERONIA homolog from *Chara braunii* was used as an outgroup. The predicted presence or absence of domains in each homolog is indicated on the right. SHRK1 and SHRK2/BSR050/SIMP2 previously characterised in *A. thaliana* are marked with an arrow. Subgroups III-A and III-B, which likely diverged in the most recent common ancestor of seed plants, are labelled. MLD - Malectin-like Domain, LRRs - Leucine-rich Repeats and Kinase - Kinase domain.

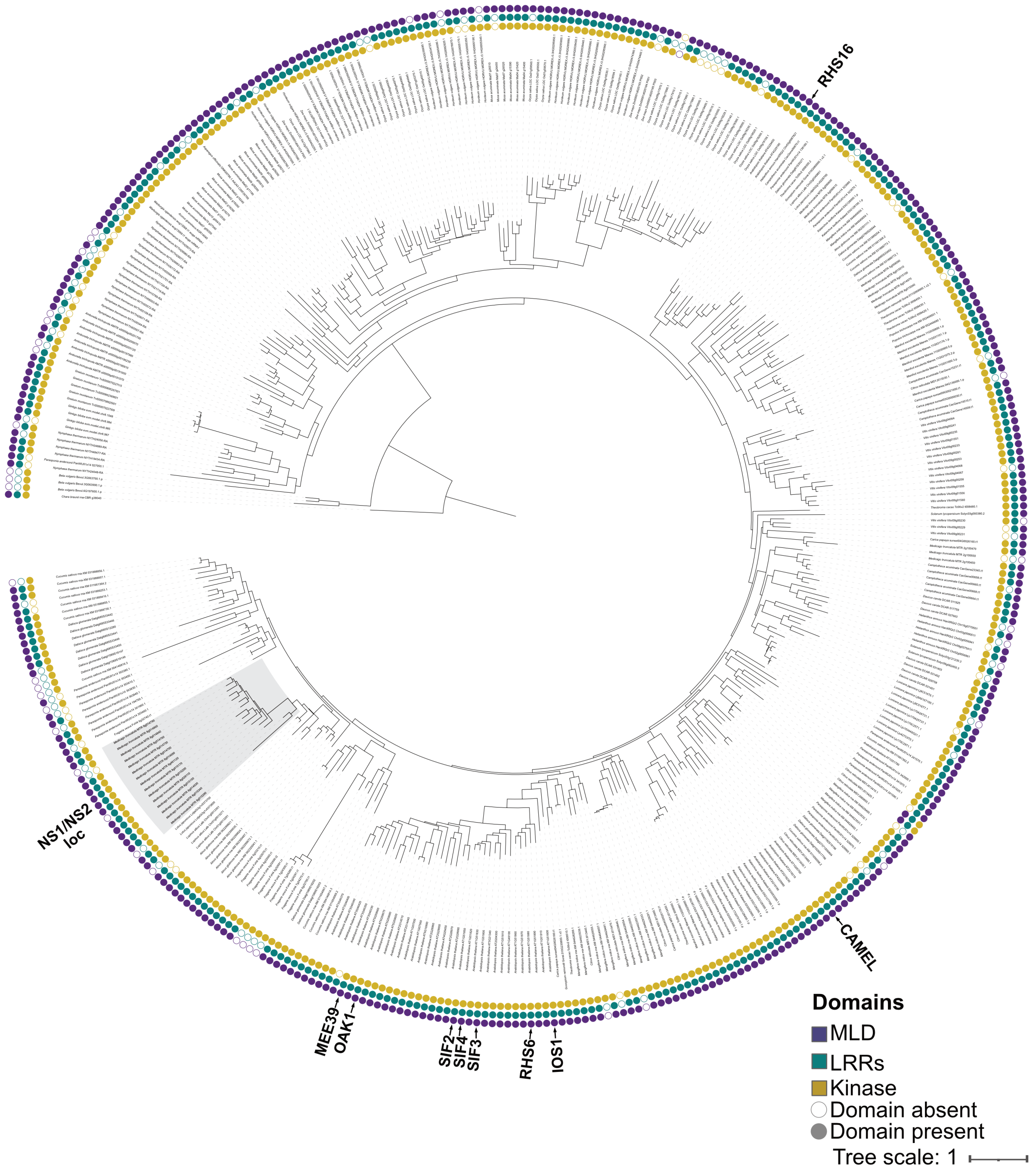


**Supplementary Figure 8: Phylogenetic tree for clade IV homologs identified in selected species.** The tree was constructed using IQtree with protein sequences assigned to clade IV and a FERONIA homolog from *Chara braunii* as an outgroup. The predicted presence and absence of MLD, LRR and Kinase domains is indicated on the outer ring of the tree. Genes previously characterised in *A. thaliana* and *Medicago truncatula* are marked with an arrow. MLD - Malectin-like Domain, LRRs - Leucine-rich Repeats and Kinase - Kinase domain.

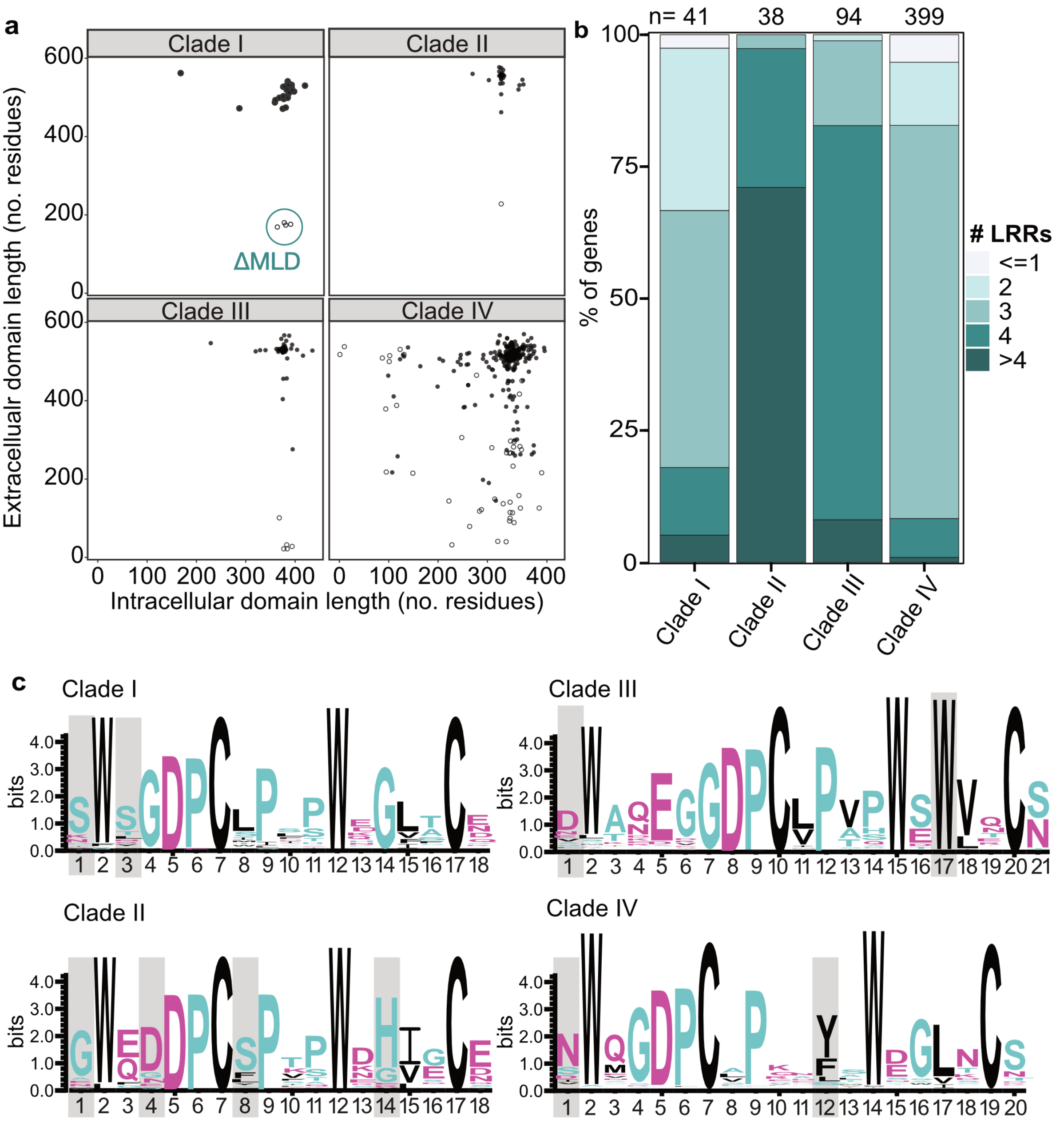


**Supplementary Figure 9: Patterns of structural variation in members of the SymRK receptor family. a -** Scatter plot showing the relationship between extracellular and intracellular domain lengths (amino acid residues) across all members of the major clades (I-IV). Proteins with canonical MLD-LRR-RLK architecture are marked with filled circles, while non-canonical variants are shown as empty circles. SymRK variants lacking the malectin-like domain are outlined (ΔMLD). **b -** Diversity in LRR numbers across the dataset. A stacked bar chart shows the frequency distribution of the number of LRR motifs per gene. The total number of genes (n) encoding LRRs is displayed at the top of the graph. **c –** Amino acid sequence logos of the ectodomains GDPC motif with flanking regions across identified major clades I-IV. Clade-specific polymorphisms are highlighted in grey. Sequence logos were generated using WebLogo (Crooks et al. 2004).

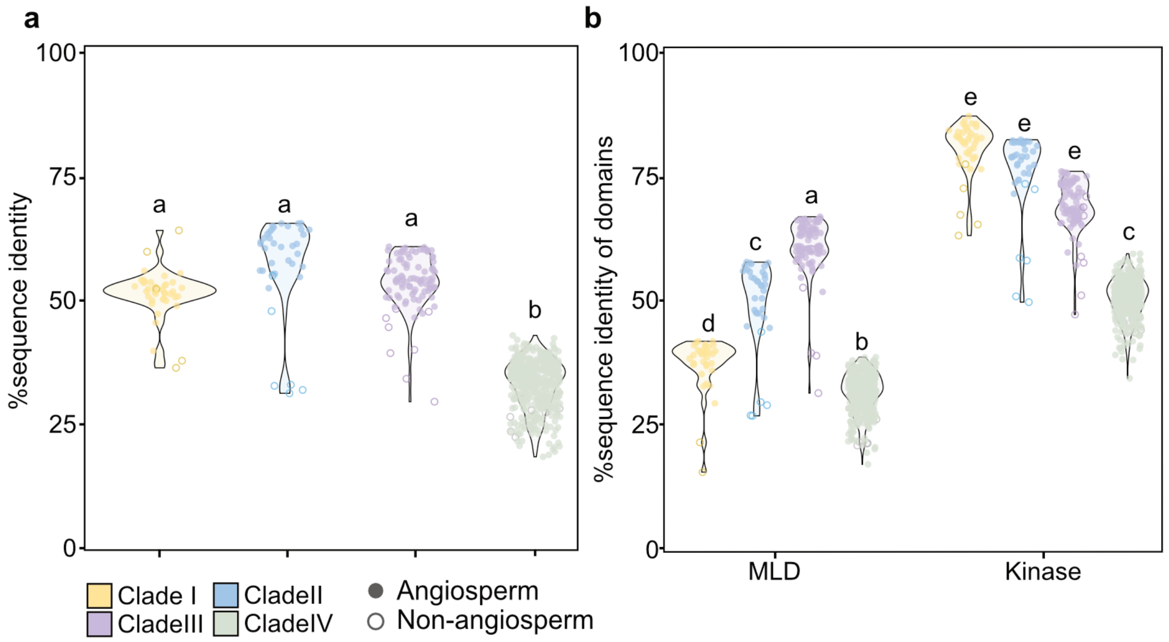


**Supplementary Figure 10: Residue-level polymorphism reveals asymmetric evolutionary patterns between extracellular and intracellular domains of SymRK homologs. a -** Dot plot showing overall sequence identity among all members of the major clades I-IV at the full-length protein level. **b -** Dot plot showing sequence identity calculated separately for the Malectin-like Domain (MLD) and the kinase domain (Kinase). In **a** and **b,** different letters indicate statistically significant differences assessed using the Kruskal-Wallis test and Dunn’s post-hoc test. In **b**, significance tests were performed across the MLD and Kinase domains together.

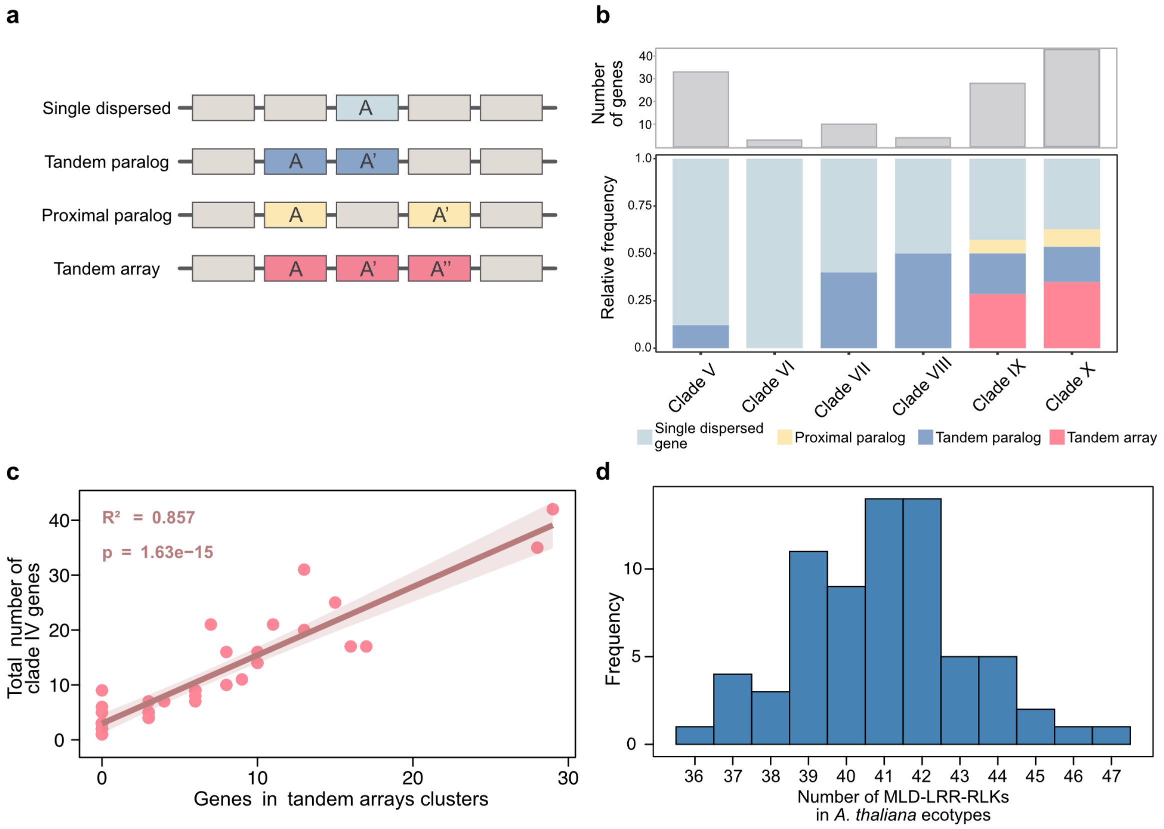


**Supplementary Figure 11: Species-specific gene duplication events drive the expansion of the MLD-LRR-RLK receptor family. a –** Schematic representation of the different classes of gene arrangement and duplications analysed in the MLD-LRR-RLKs. **b -** Top: Total number of genes assigned to each of the V-X clades. Bottom: Relative frequencies of clade-specific genes occurring either as single genes or with additional nearby copies classified as proximal paralogs, tandem paralogs or tandem arrays as in a). **c -** Scatter plot and linear model showing the relationship between the total clade IV members and the number of genes in tandem arrays across species with clade IV genes. **d –** Frequency distribution of the total number of MLD-LRR-RKs identified across 69 *A. thaliana* ecotypes.

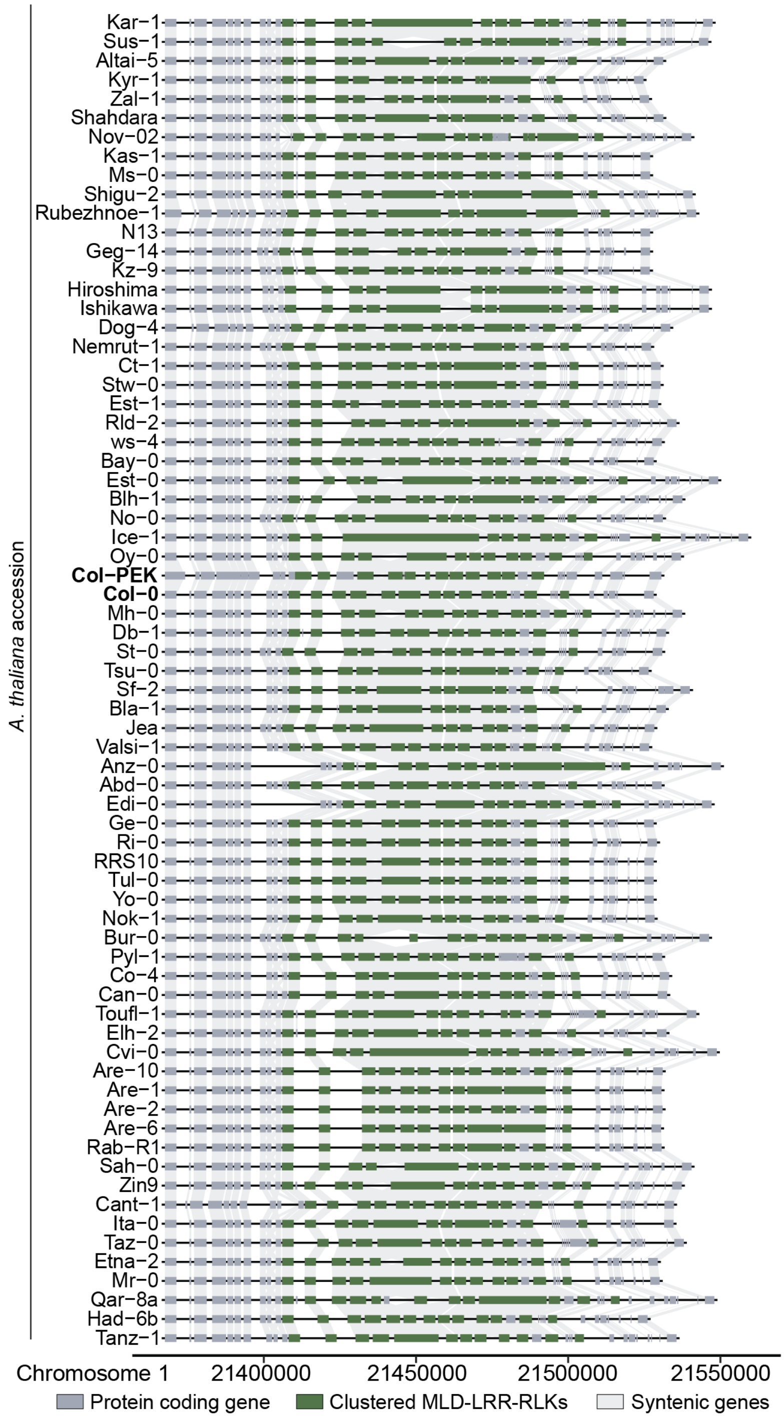


**Supplementary Figure 12: Local synteny of clustered clade IV members on chromosome 1 in *Arabidopsis thaliana*.** Syntenic diversity of SymRK receptor family members clustered on chromosome 1 across 69 *A. thaliana* accessions. Genes encoding canonical MLD-LRR-RLKs are shown in dark green, non-canonical variants in light green and flanking protein-coding genes within ± 10 kb in grey. Accession names are listed on the left. Col-0 and Col-0 PEK (Hou et al. 2022) (with improved Col-0 genome assembly) were used as reference genomes for the synteny map. Numbers on the x-axis indicate genomic coordinates along chromosome 1.

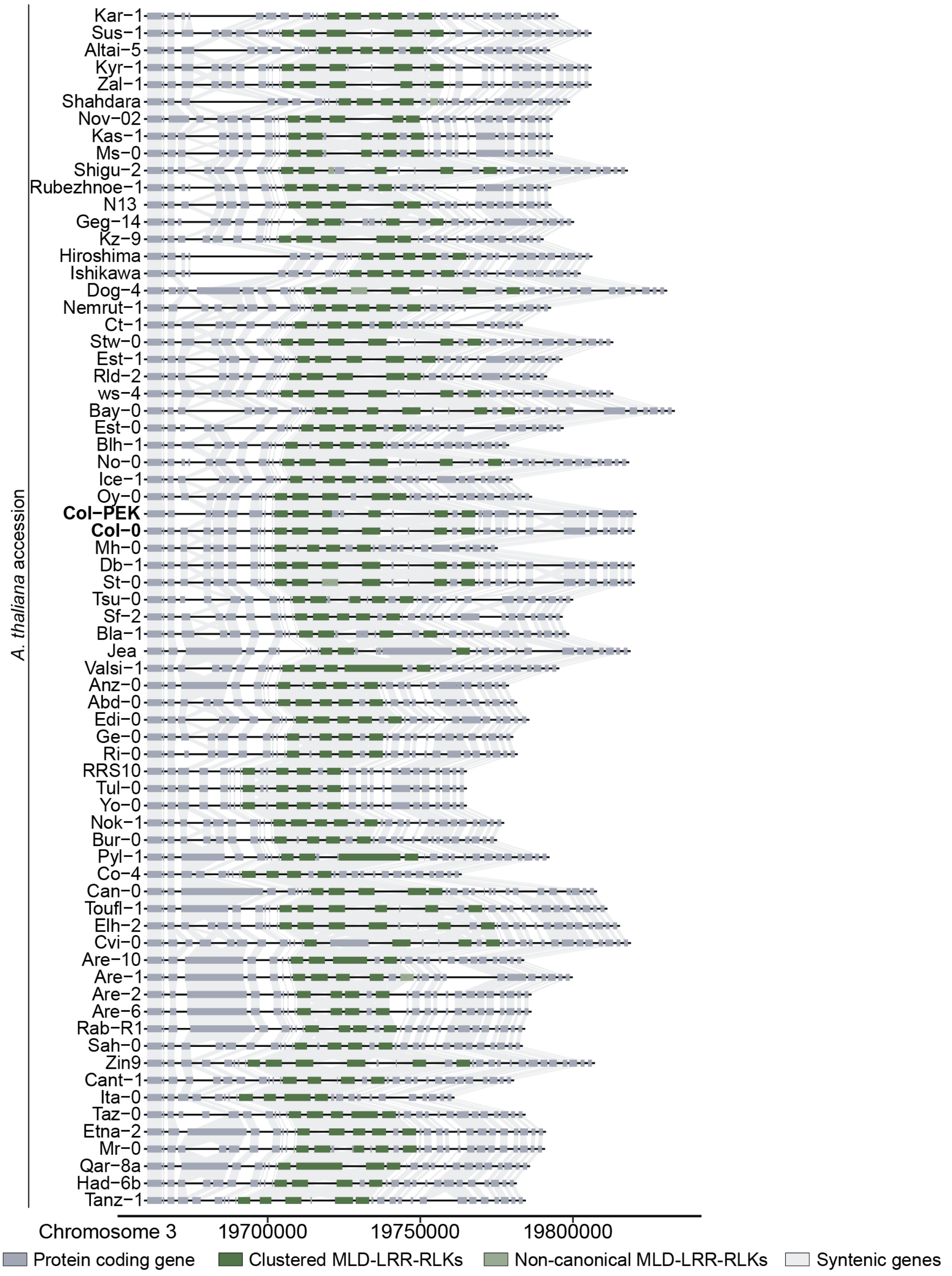


**Supplementary Figure 13: Local synteny of clustered clade IV members on chromosome 3 in *Arabidopsis thaliana*.** Syntenic diversity of SymRK receptor family members clustered on chromosome 3 across 69 *A. thaliana* accessions. Genes encoding canonical MLD-LRR-RLKs are shown in dark green, non-canonical variants in light green and flanking protein-coding genes within ± 10 kb in grey. Accession names are listed on the left. Col-0 and Col-0 PEK (Hou et al. 2022) (with improved Col-0 genome assembly) were used as reference genomes for the synteny map. Numbers on the x-axis indicate genomic coordinates along chromosome 3.

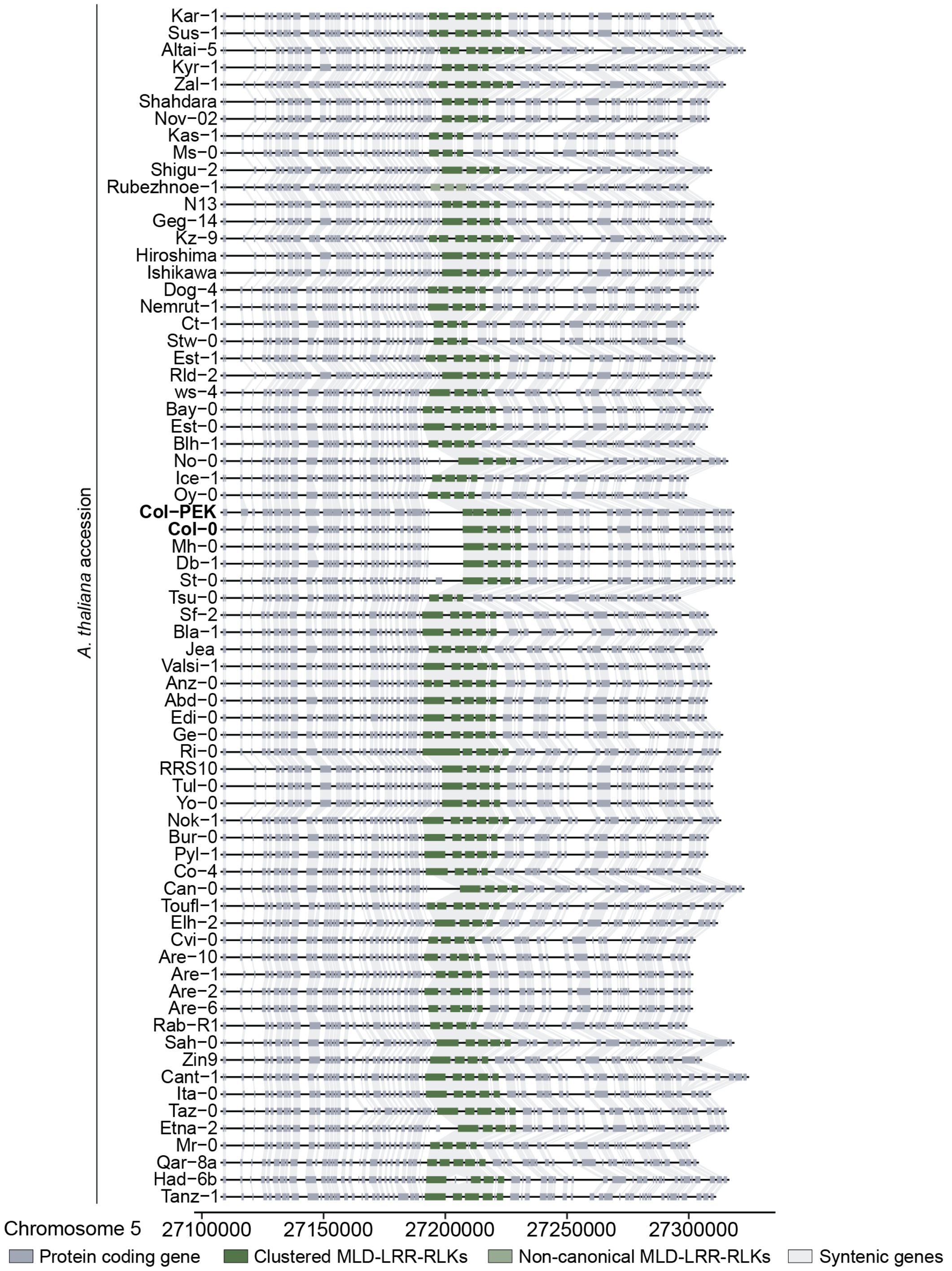


**Supplementary Figure 14: Local synteny of clustered clade IV members on chromosome 5 in *Arabidopsis thaliana*.** Syntenic diversity of SymRK receptor family members clustered on chromosome 5 across 69 *A. thaliana* accessions. Genes encoding canonical MLD-LRR-RLKs are shown in dark green, non-canonical variants in light green and flanking protein-coding genes within ± 10 kb in grey. Accession names are listed on the left. Col-0 and Col-0 PEK (Hou et al. 2022) (with improved Col-0 genome assembly) were used as reference genomes for the synteny map. Numbers on the x-axis indicate genomic coordinates along chromosome 5.

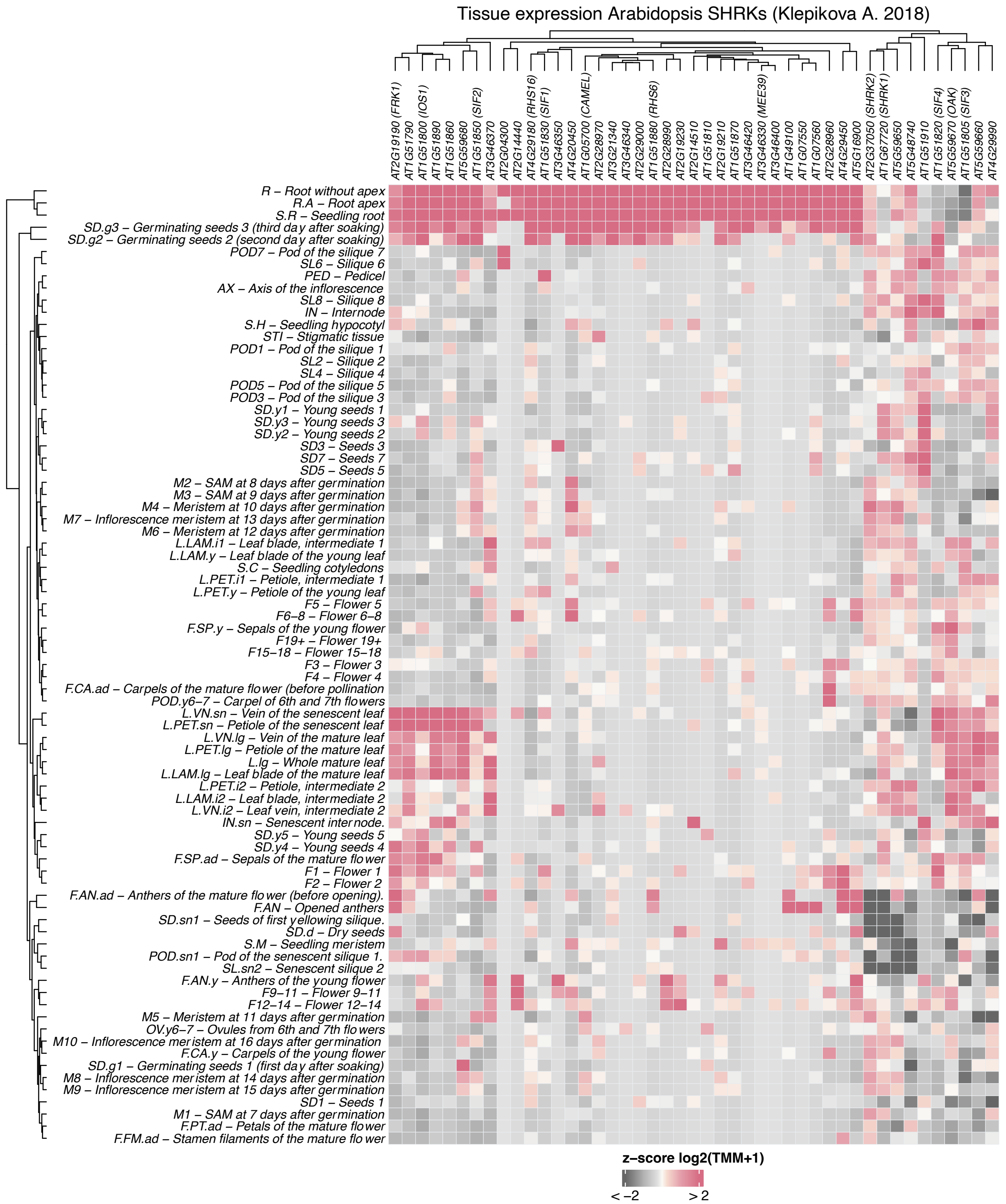


**Supplementary Figure 15: Most of *Arabidopsis thaliana MLD-LRR-RLKs* are expressed in the root.** The heat map depicts transcriptional expression patterns of *A. thaliana MLD-LRR-RLKs* across multiple tissues, organs and developmental stages, listed row-wise on the left of the heat map. Expression data, derived from (Klepikova et al. 2016), are shown as normalised gene counts, with transcript abundance represented by shades of grey and pink. Genes were hierarchically clustered based on similarities in their co-expression profiles. Gene IDs are shown at the top of the heat map, accompanied by common names where available.

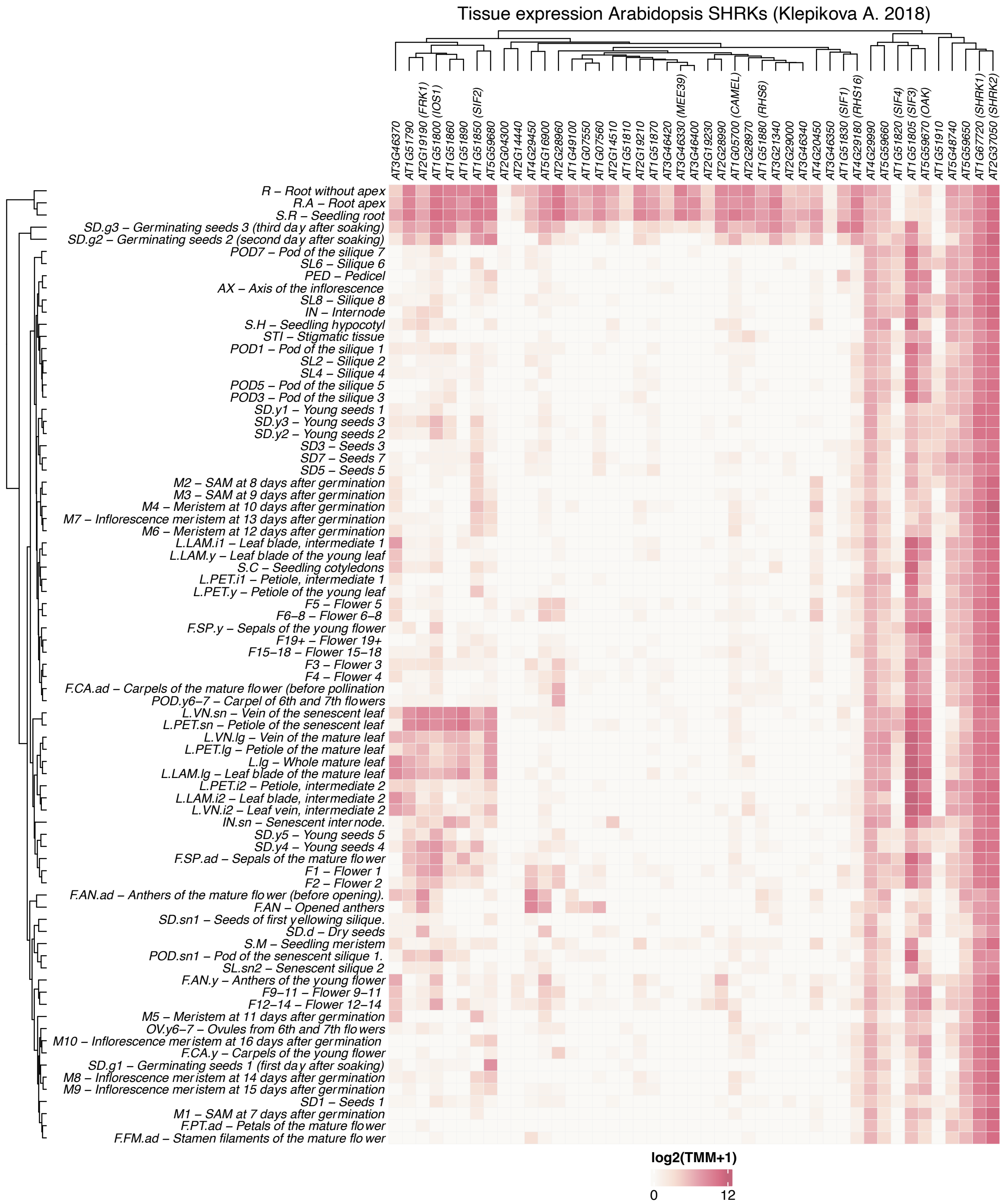


**Supplementary Figure 16: *MLD-LRR-RLKs* generally exhibit low expression levels across different tissues in *Arabidopsis thaliana*.** The heat map depicts transcriptional expression patterns of *A. thaliana MLD-LRR-RLKs* across multiple tissues, organs and developmental stages, listed row-wise on the left of the heat map. Expression data, derived from (Klepikova et al. 2016), are shown as total gene counts, with transcript abundance represented by shades of ivory and pink. Genes were hierarchically clustered based on similarities in their co-expression profiles. Gene IDs are shown at the top of the heat map, accompanied by common names where available.

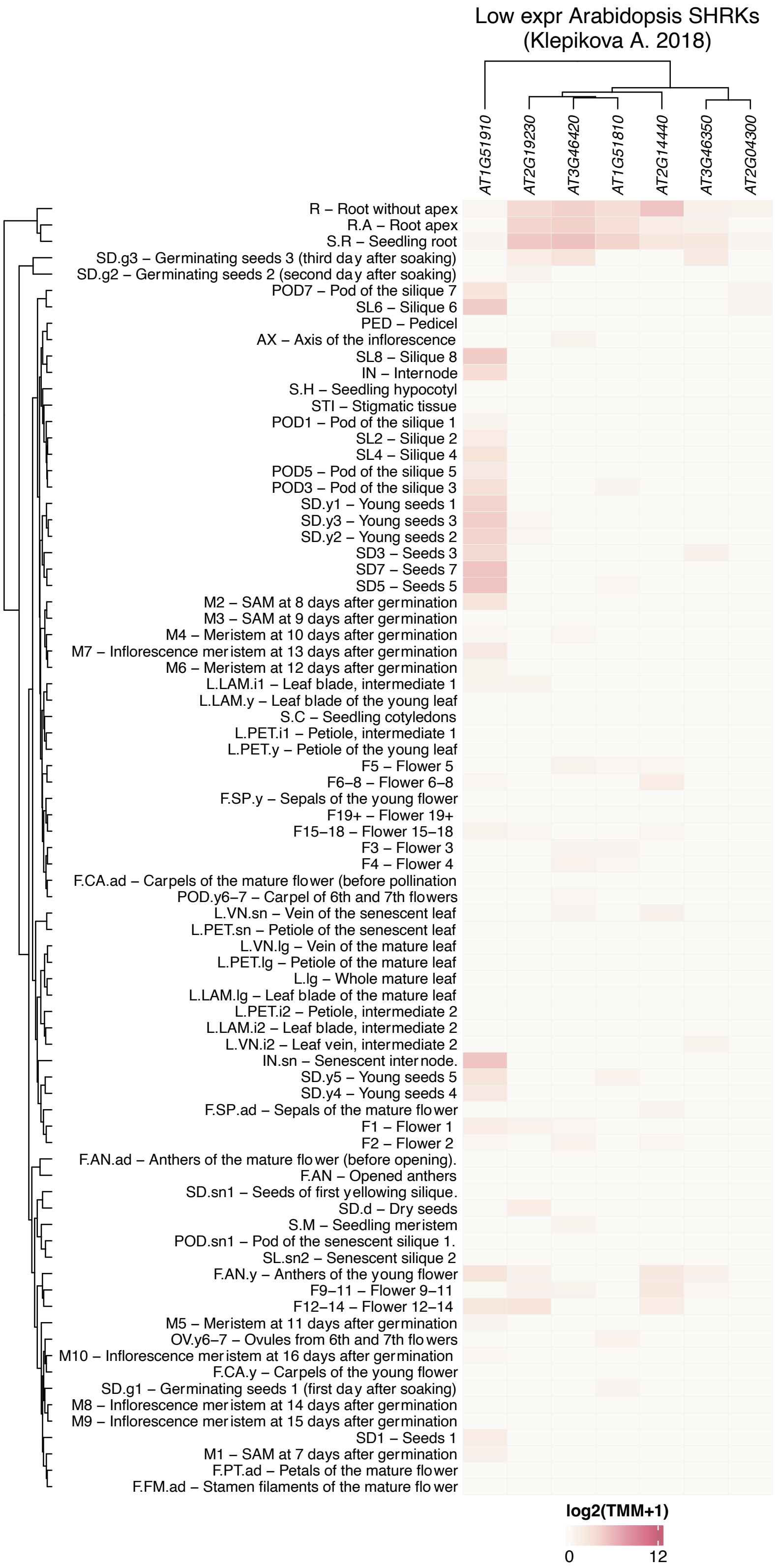


**Supplementary Figure 17: A subset of *MLD-LRR-RLKs* exhibits minimal expression levels across different tissues in *Arabidopsis thaliana*.** The heat map depicts transcriptional expression patterns of *A. thaliana MLD-LRR-RLKs* across multiple tissues, organs and developmental stages, listed row-wise on the left of the heat map. Expression data, derived from (Klepikova et al. 2016), are shown as total gene counts, with transcript abundance represented by shades of ivory and pink.

Genes were hierarchically clustered based on similarities in their co-expression profiles. Gene IDs are shown at the top of the heat map.

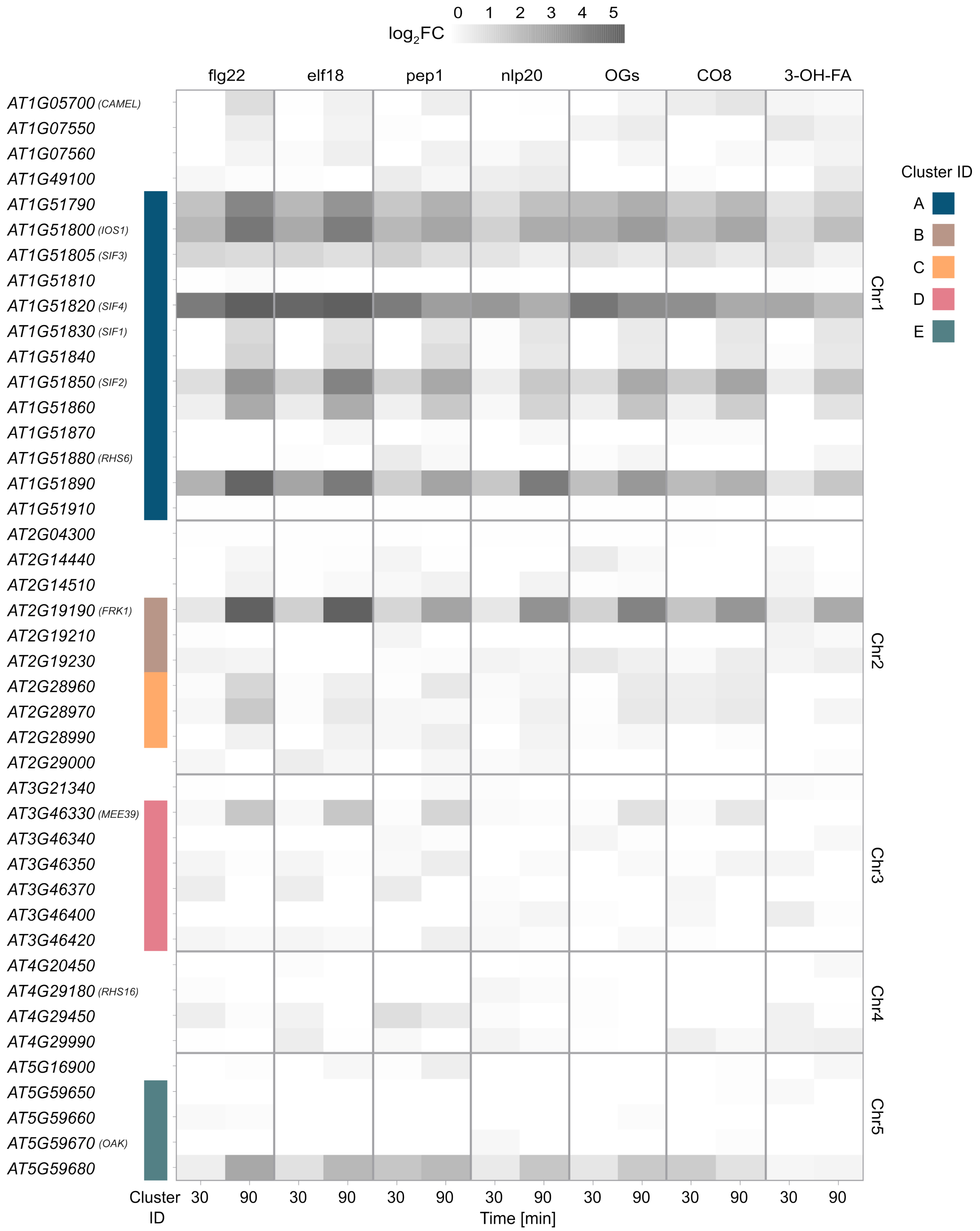


**Supplementary Figure 18: Transcriptional responses of *Arabidopsis thaliana* clade IV genes to elicitor treatments.** Heat map showing the expression changes for clade IV members in response to various elicitors (Bjornson et al. 2021) at two time points (30 and 90 minutes after exposure) in seedlings. Gene IDs are listed on the left. Each cell represents the $\log_2$ fold change in transcript abundance relative to untreated controls. The colour scale bar is shown above the heat map. flg22: 22-amino-acid epitope derived from bacterial flagellin; elf18: 18-amino-acid epitope derived from bacterial elongation factor Tu; pep1: 23-amino-acid peptide released as a DAMP upon cellular damage; nlp20: 20-amino-acid peptide derived from bacterial, oomycete and fungal NECROSIS AND ETHYLENE-INDUCING PEPTIDE 1 - LIKE PROTEINS; CO8: chitooctaose, an octamer fragment of fungal cell walls; OGs: oligogalacturonides, derived from the plant cell wall; 3-OH-FA: bacterial hydroxylated fatty acid.

### References:


Bjornson M, Pimprikar P, Nürnberger T, Zipfel C. 2021. The transcriptional landscape of *Arabidopsis thaliana* pattern-triggered immunity. Nat Plants. 7**:** 579-586. 10.1038/s41477-021-00874-5.

Crooks GE, Hon G, Chandonia JM, Brenner SE. 2004. Weblogo: A sequence logo generator. Genome Res. 14**:** 1188-1190. 10.1101/gr.849004.

Hou X, Wang D, Cheng Z, Wang Y, Jiao Y. 2022. A near-complete assembly of an *Arabidopsis thaliana* genome. Mol Plant. 15**:** 1247-1250. 10.1016/j.molp.2022.05.014.

Klepikova AV, Kasianov AS, Gerasimov ES, Logacheva MD, Penin AA. 2016. A high resolution map of the *Arabidopsis thaliana* developmental transcriptome based on rna-seq profiling. The Plant Journal. 88**:** 1058-1070. https://doi.org/10.1111/tpj.13312.
